# Molecular mechanisms underlying p62-dependent secretion of the Alzheimer-associated ubiquitin variant, UBB^+1^

**DOI:** 10.1101/2024.12.31.630908

**Authors:** Ajay R. Wagh, Michael H. Glickman

**Author notes:** Corresponding author: The Faculty of Biology #438, Technion Israel Institute of Technology, Haifa, 3200003, Israel.

## Abstract

UBB⁺¹, a ubiquitin variant protein resulting from a frameshift in the ubiquitin-B gene, is a pathological hallmark of Alzheimer disease (AD). At the cellular level, UBB⁺¹ disrupts the ubiquitin-proteasome system while inducing autophagy. Notably, UBB⁺¹ itself is secreted via autophagosome-like vesicles. Here, we demonstrate that UBB⁺¹ can be removed from the cell by degradative and secretory autophagy. SQSTM1/p62 functions as a pivotal ubiquitin receptor for UBB⁺¹, recognizing its ubiquitin domain and facilitating loading into autophagosomes. Oligomerization of SQSTM1/p62 was critical to isolate UBB⁺¹ in bodies preventing its aggregation. Intriguingly, both gain- and loss-of-function SQSTM1/p62 suppressed UBB⁺¹ secretion, causing intracellular retention: SQSTM1/p62 knockout led to UBB⁺¹ accumulation in insoluble aggregates, while its overexpression promoted the formation of p62-UBB⁺¹ bodies. We further identified distinct roles for SNARE-mediated membrane fusion in secretory autophagy of UBB⁺¹. Specifically, the R-SNARE SEC22B and the Q-SNAREs Syntaxin-4 (STX4) and SNAP23 participated in UBB⁺¹ exocytosis. Disruption of SEC22B impaired the fusion of UBB⁺¹-containing autophagosomes with the plasma membrane, reducing UBB⁺¹ secretion without affecting its intracellular turnover. Inhibition of lysosomes partially stabilized UBB⁺¹ indicating that degradation and secretion are complementary processes that determine the fate of UBB^+1^. This study elucidates the dual roles of autophagy in managing neurotoxic proteins, highlighting SQSTM1/p62 as a key mediator of UBB⁺¹ trafficking and secretion. Although ubiquitin typically acts as a degradation signal, our findings reveal a rare instance of a ubiquitin-related protein driving secretory autophagy. These findings advance our understanding of cellular mechanisms underlying the clearance of misfolded proteins in neurodegenerative diseases.

**Significance:** To maintain health, cells must remove toxic proteins, typically by ubiquitin-dependent degradation. Neurons are particularly sensitive since they are not dividing and cannot replace damaged cells. This study suggests that autophagy, a pathway for degrading proteins, can also secrete harmful proteins, highlighting a new pathway for tackling neurodegenerative diseases. Specifically, UBB^+1^, a variant of the ubiquitin protein linked to Alzheimer’s disease, is recognized by a central autophagy component, p62, packaged into vesicles and secreted from cells. When p62 is absent, UBB^+1^ accumulates inside cells, which can be harmful. This work identifies UBB⁺¹ as a novel cargo for secretory autophagy, extending the understanding of how cells may handle proteotoxic stress beyond classical degradation pathways.

## Introduction

Autophagy is traditionally viewed as a degradative process (1), but growing evidence has revealed its role in vesicle production (2), the secretion of proteins such as pro-inflammatory cytokines (3), or lysozyme release (4). These processes, collectively termed autophagy-associated secretion, or secretory autophagy, define an autophagy pathway that plays an active role in maintaining cellular homeostasis in both healthy and disease states (2-4). Unlike conventional degradative autophagy, where cytosolic cargoes are broken down, secretory autophagy involves the transport of these cargoes within autophagosomes to the extracellular environment through exocytosis (5). This secretory autophagy machinery has been implicated in the exocytosis of various elements, including cytokines (such as IL-1β, IL-6, IL-18, and TNF-α), aggregated proteins, organelles, and microbial components (5, 6). An important role of degradative autophagy, by contrast, is the removal of ubiquitinated cargo such as ubiquitinated proteins, ubiquitinated aggregates (aggrephagy), and ubiquitinated organelles (e.g., mitophagy) (7, 8). While many studies have explored autophagy-associated protein secretion/release, few have focused on the recruitment of cargo into secretory autophagosomes.

Several dedicated ubiquitin receptors facilitate the loading of cargo into autophagosomes, serving as bridges between ubiquitinated cargo and the isolation membrane (9). These receptors possess specific domains: a ubiquitin recognition element on one side and an LC3-interacting region (LIR) on the other. One classical autophagy receptor is p62/sequestosome 1 (SQSTM1), which plays a crucial role in clearing ubiquitinated proteins (10, 11). SQSTM1/p62 abnormal expression or SQSTM1gene mutation is tightly associated with various diseases including AD. SQSTM1/p62 has a multidomain structure tailored for this function: its N-terminal PB1 domain mediates self-oligomerization, a central LIR motif targets it to the phagophore, and a C-terminal ubiquitin-associated (UBA) domain facilitates ubiquitin binding (9, 12, 13). Through its UBA domain, SQSTM1/p62 noncovalently binds monoubiquitin and Lys48- or Lys63-linked polyubiquitin chains (14-16). Notably, proteins with UBA domains generally exhibit a stronger preference for binding polyubiquitin chains over monoubiquitin (17, 18).

Misfolded protein aggregation in both intracellular and extracellular environments is a key factor in the development of Alzheimer disease (AD). Neurons rely on autophagy for the removal of defective proteins to maintain neuronal health (19-22). The primary pathological hallmarks of AD are the formation of amyloid plaques, which are primarily composed of aggregated amyloid-β (Aβ) peptides, and intracellular neurofibrillary tangles containing hyper-phosphorylated tau (23-25). UBB^+1^ is a non-hereditary mutation of ubiquitin that results from a transcription error leading to a dysfunctional ubiquitin. UBB^+1^ is associated with AD by accumulating in both extracellular plaques and intracellular tangles of patients (26). UBB⁺¹ arises from transcriptional frameshift errors in the ubiquitin B gene and accumulates robustly in neurofibrillary tangles and amyloid plaques of advanced Alzheimer’s disease patients (26-28). A transgenic mouse model further confirmed UBB⁺¹ deposition in hippocampal and brainstem regions associated with AD-like phenotypes, including memory deficits and proteostasis dysfunction (29, 30). Human post-mortem studies have also reported UBB⁺¹ accumulation as early as Braak stages III–IV (25), highlighting its progressive buildup. Although the precise ratio of UBB⁺¹ to wild-type ubiquitin in vivo is difficult to measure, its persistent presence across disease stages points to its biological significance.

This dual intra- and extra-cellular localization raises questions about the pathway by which UBB^+1^ exits the cell. Indeed, in our previous study, we investigated this pathway and demonstrated that autophagy components play a role in the unconventional secretion of UBB^+1^ (31). Unconventional secretion involves the packaging of cytosolic proteins into vesicles which ultimately fuse to the plasma membrane to release their contents into the extracellular space (32, 33). This process is mediated by SNARE proteins, which catalyze membrane fusion. Among the 38 known mammalian SNARE proteins, each is selectively localized to specific cellular membranes, thereby defining distinct trafficking pathways (34). SNARE proteins are classified based on the central amino acid residues in their SNARE motifs: Q-SNAREs for glutamine and R-SNAREs for arginine. A functional SNARE complex required for membrane fusion consists of a parallel four-helix bundle formed by one Qa-, Qb-, Qc-, and R-SNARE. Specific sets of these proteins catalyze distinct trafficking pathways.

In this study, we show that autophagy-mediated secretion of UBB^+1^ is dependent on SQSTM1/p62. Knockout (KO) of p62 in HeLa cells significantly disrupts UBB^+1^ recruitment to SEC22B-containing secretory autophagosome-like vesicles, resulting in the formation of intracellular aggregates and a marked inhibition of its secretion. We further elucidate the participation of the SNARE proteins, SEC22B, STX4 and SNAP23 in UBB^+1^ exocytosis. These findings provide novel perspectives on the cellular mechanisms involved in the removal of misfolded proteins that can cause significant damage to the cell and lead to neurodegenerative diseases.

## Results

### Secretion of UBB^+1^ shares properties with secretory autophagy

In secretory autophagy, SEC22B is one of the factors responsible for mediating fusion with the plasma membrane and plays an essential role in the clearance of an autophagic substrate (35, 36). We generated *sec22b* KO HeLa cells (**Fig. S1A)** and confirmed that loss of SEC22B impairs autophagic flux, resulting in autophagosome accumulation. Expression of UBB^+1^ induces LC3B-II, which can be visualized as puncta (**Fig. 1A and B and Fig. S1B**). In *sec22b* KO cells, UBB⁺¹ secretion is suppressed, and colocalization with LC3B is more readily observed, consistent with impaired autophagic flux and blocked vesicle clearance (**Fig. 1B and Fig. S1C**).

**Figure 1.**
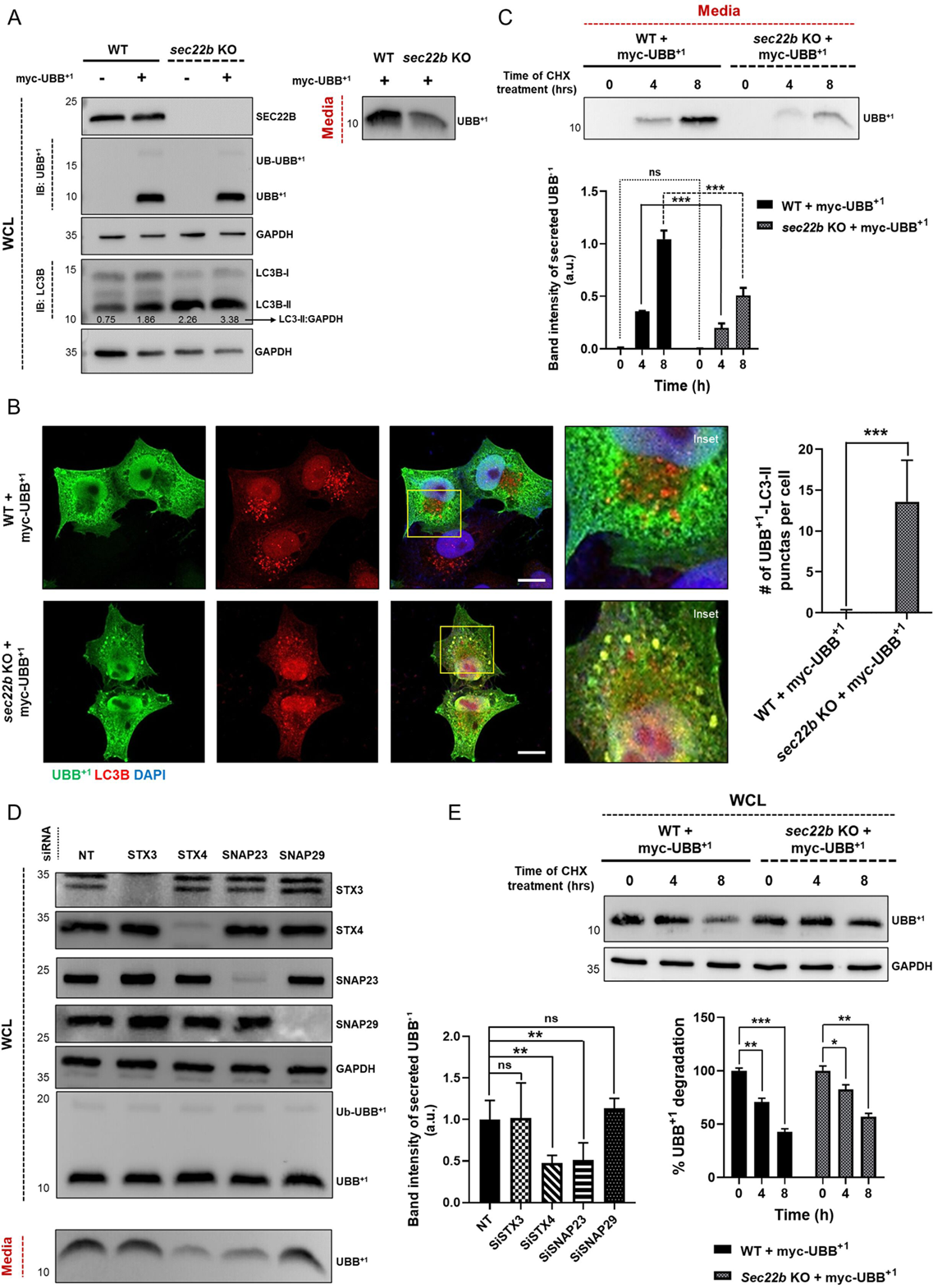
Q- and R-SNAREs involved in UBB^+1^ secretion. **(A)** Wild-type (WT) and *sec22b* knockout (KO) HeLa cells were transiently transfected with a MYC-UBB⁺¹ expressing plasmid or treated with transfection reagent alone (mock control). After 24 h, cells were incubated with fresh DMEM for an additional 16 h. Conditioned media and whole-cell lysates (WCL) were collected, resolved by 15% SDS-PAGE, and immunoblotted with specific antibodies. UBB⁺¹ and LC3 were detected to assess secretion and autophagy induction, respectively, GAPDH was used as a loading control for WCL. Secreted UBB⁺¹ in the media was detected using anti-UBB^+1^ antibody. Equal volumes of media were collected from cultures of comparable cell numbers across all conditions to ensure consistency in secretion analysis. Band intensities of LC3-II were quantified relative to GAPDH and are indicated below the blots. **(B)** Intracellular distribution of UBB⁺¹ (Green) and LC3 (Red) was visualized by confocal immunofluorescence microscopy using specific antibodies. Nuclei were counterstained with DAPI (blue). In wild-type HeLa cells expressing MYC-UBB⁺¹, LC3-positive autophagic vesicles primarily accumulated in the juxtanuclear region with limited colocalization with UBB⁺¹ puncta. In contrast, *sec22b* KO cells displayed prominent colocalization of UBB⁺¹ with LC3-positive autophagosomes, consistent with impaired autophagosome maturation and reduced vesicle clearance. Scale bar: 5 μm. Quantification of UBB⁺¹–LC3 colocalized puncta was performed by analyzing 100 randomly selected cells per condition from three independent experiments. Bar graphs show mean ± SD. Statistical significance was determined using an unpaired two-tailed Student’s t-test; *, **, and *** denote p < 0.05, 0.01, and 0.001, respectively; ns = non-significant. **(C)** WT and *sec22b* KO HeLa cells expressing MYC-UBB⁺¹ were incubated with 100 μg/ml cycloheximide (CHX) for the indicated time periods. Conditioned media were collected in equal volumes from comparable cell numbers across all experimental conditions and immunoblotted for UBB⁺¹ (top). Densitometric analysis of UBB⁺¹ secretion (bottom) was performed by quantifying the UBB⁺¹ band intensity. For each time point, UBB⁺¹ levels in *sec22b* KO cells were compared to WT cells. Statistical comparisons between WT and *sec22b* KO are indicated in the graph: comparisons at 0 h are shown by round dotted lines, at 4 h by solid lines, and at 8 h by square dotted lines. Data represents the mean ± SD from three independent experiments. Statistical significance was determined using an unpaired two-tailed t-test. *, **, and *** denote p < 0.05, 0.01, and 0.001, respectively; ns = non-significant. In the bar graph, *sec22b* KO data are shown in [gray], and WT data are shown in [black]. **(D)** HeLa cells expressing MYC-UBB⁺¹ were transfected with 25 nM DsiRNAs targeting STX3, STX4, SNAP23, or SNAP29, or with a non-targeting (NT) control (see Table 1 for sequences). Forty-eight hours after transfection, cells were incubated in fresh media for 16 h. Conditioned media and corresponding whole-cell lysates (WCLs) were collected and analyzed by SDS-PAGE followed by immunoblotting. Seven blots show: STX3, STX4, SNAP23, and SNAP29 knockdown efficiency, GAPDH as the WCL loading control, UBB⁺¹ levels in WCLs, and secreted UBB⁺¹ in the media. Conditioned media were collected in equal volumes from comparable cell numbers across all experimental conditions and immunoblotted for UBB⁺¹. STX4 and SNAP23 knockdown significantly reduced UBB⁺¹ secretion, whereas STX3 and SNAP29 knockdown had no notable effect. Data are representative of three independent experiments; quantification and statistical analysis were performed using one-way ANOVA. *, **, and *** denote p < 0.05, 0.01, and 0.001, respectively; ns = non-significant. **(E)** WT and *sec22b* KO HeLa cells expressing MYC-UBB⁺¹ were incubated with 100 μg/ml cycloheximide (CHX) for the indicated periods. The cells were then harvested, and WCL immunoblotted for UBB⁺¹ (top). Densitometric analysis of the UBB⁺¹ relative to untreated cells (normalized to GAPDH content) was averaged from three independent experiments (bottom). The error bars represent the means ± SD. Statistical significance was determined using an unpaired two-tailed t-test. For all statistical tests *, **, ***, *p* < 0.05, 0.01, and 0.001, respectively, ns = non-significant.

To confirm whether these autophagosomes are involved in UBB^+1^ secretion, we utilized *sec22b* KO cells overexpressing UBB^+1^ and evaluated its release to the growth media (**Fig. S1D**). Treatment of UBB^+1^-expressing *sec22b* KO cells with cycloheximide (CHX), a protein synthesis inhibitor, showed that its secretion was significantly reduced compared to wild-type cells (**Fig. 1C**). To rule out the possibility that secretion was caused by cytotoxicity or apoptosis under CHX treatment or other experimental conditions, we assessed cleaved caspase-3 levels by Western blot. No cleaved caspase-3 was detected in WT or *sec22b* KO cells expressing UBB⁺¹ with or without CHX or CHX + Bafilomycin A1. In contrast, a strong cleaved caspase-3 band was observed in cells treated with Venetoclax (ABT-199), a known inducer of intrinsic apoptosis (37), confirming the effectiveness of the assay (**Fig. S1G**). These results demonstrate that UBB⁺¹ secretion under these conditions is not due to apoptotic cell death.

At the plasma membrane, secretory autophagosomes containing SEC22B, an R-SNARE, fuse with Q-SNAREs such as STX3 and STX4 (38, 39). As shown in **Figure 1D**, knockdown of STX4 by dicer-substrate small interfering RNA (DsiRNA) reduced UBB^+1^ secretion. Conversely, knockdown of STX3 had no effect on UBB^+1^ secretion (**Fig. 1D**). To further clarify the role of SNARE machinery, we performed knockdown experiments targeting additional Q-SNARE proteins SNAP23 and SNAP29. Knockdown of SNAP23 significantly impaired secretion of UBB⁺¹, similarly to STX4 knockdown, while SNAP29 knockdown had no significant effect compared to cells treated with non-targeting (NT) siRNA (**Fig. 1D**). These findings indicate that SNAP23, together with STX4, are Q-SNAREs participating in fusion of UBB⁺¹-containing secretory autophagosomes with the plasma membrane. Alternatively, in degradative autophagy, STX17 facilitates autophagosome fusion with the lysosome (40-42). Knockdown of STX17 had no significant effect on UBB^+1^ secretion (**Fig. S1E**), indicating either that STX17 is redundant, or that degradative autophagy is not implicated in the secretion of UBB^+1^. Similarly, we did not find evidence for exosome-mediated secretion of UBB⁺¹, as knockdown of TSG101, a key regulator of exosome biogenesis, did not reduce UBB⁺¹ secretion (**Fig. S1F**). Consistent with this, we have previously shown that ALIX knockout HeLa cells also secrete UBB⁺¹ efficiently, further supporting that exosome pathways are not involved in UBB⁺¹ secretion (31).

To investigate the relationship between secretion and intracellular turnover dynamics, we expressed UBB^+1^ in HeLa cells and treated them with CHX. Intracellular UBB^+1^ levels decreased continuously over 4 to 8 h of CHX treatment, indicating a biological half-life of roughly eight hours (**Fig. 1E**). Nevertheless, the intracellular turnover of UBB⁺¹ persisted despite the absence of SEC22B, while its secretion was significantly impaired under the same conditions (**Fig. 1C**). Taken together, these results highlight the roles of SEC22B and STX4/SNAP23 in the secretory autophagy machinery for UBB⁺¹ secretion, while an alternative mechanism is likely operating to clear the intracellular buildup of UBB⁺¹.

### Secretion and degradation of UBB^+1^ both account for its intracellular turnover dynamics

To determine if intracellular degradation contributes to UBB^+1^ turnover, its cellular half-life was evaluated upon addition of lysosomal (Bafilomycin) or proteasomal (Velcade) inhibitors and CHX treatment. As shown in **Figure 2A**, intracellular UBB⁺¹ levels significantly decreased during CHX treatment, irrespective of inhibitor treatment. CHX and Velcade efficiency were confirmed as shown in **Figure S2A and S2B**. These turnover dynamics indicate that intracellular degradation alone cannot fully account for the clearance of UBB⁺¹ from the cells. In contrast, the effect of intracellular proteolysis inhibitors on extracellular UBB⁺¹ levels was minimal (**Fig. 2B**), suggesting that secretion serves as a compensatory mechanism when degradation pathways are impaired. In *sec22b*-deficient cells, where UBB⁺¹ secretion is significantly impaired but not completely abolished, treatment with Bafilomycin further enhanced extracellular UBB⁺¹ levels (**Fig. 2C and 2D**). This suggests that when lysosomal degradation is blocked, UBB⁺¹ may be redirected toward alternative vesicular pathways for secretion, even in the absence of SEC22B. In parallel, the intracellular pool of UBB^+1^ was partially stabilized (**Fig. 2E and 2F**). These data support our earlier observation (31) that UBB^+1^ degradation and secretion are complementary processes that together determine the fate of UBB^+1^. We conclude that UBB^+1^-containing autophagosome-like vesicles can be targeted either to the lysosome or to the plasma membrane, effectively distributing their cargo between these two destinations. This observation prompted further investigation into the mechanisms of how UBB^+1^ is packaged into these vesicles.

**Figure 2.**
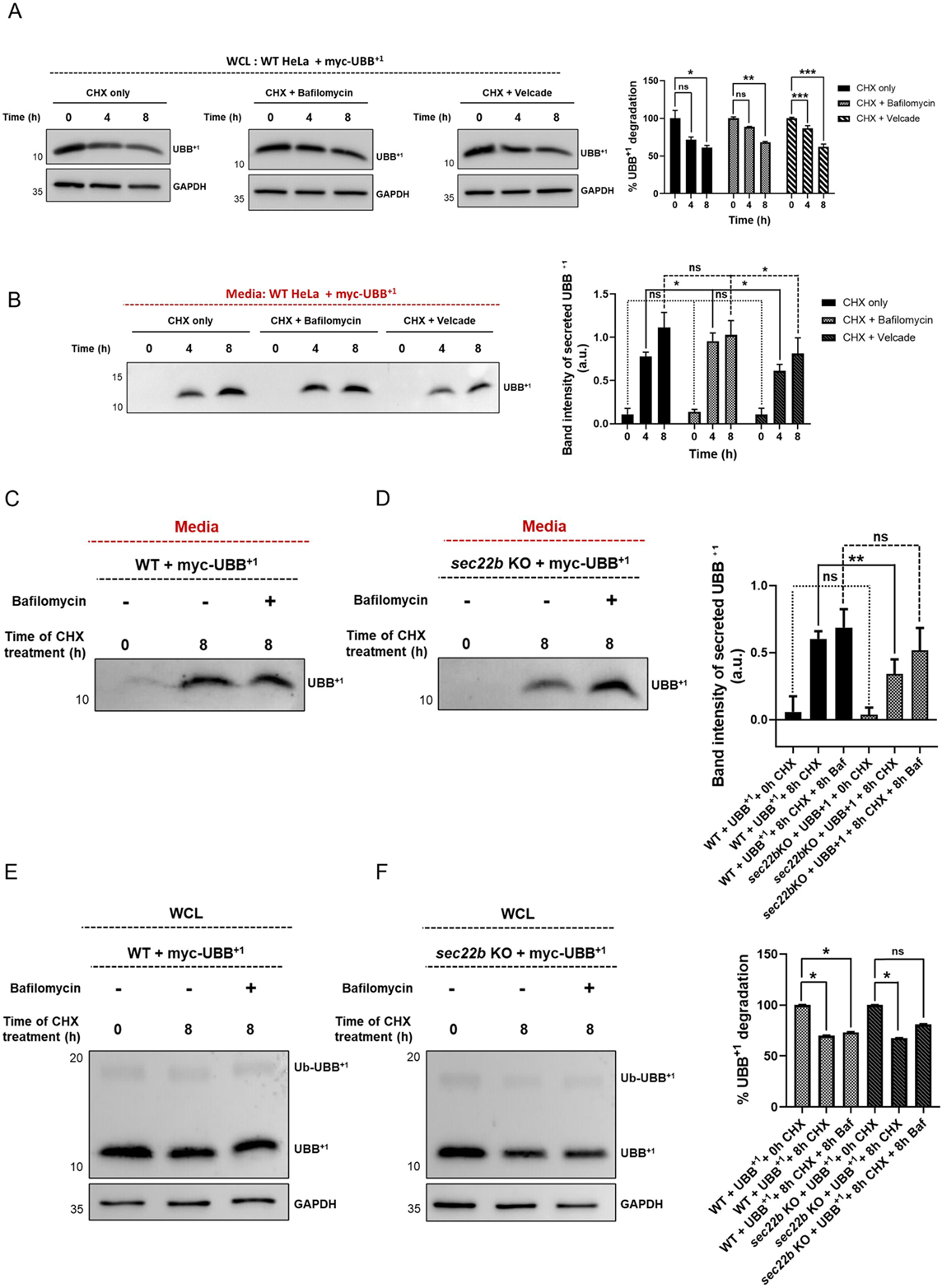
Turnover and Secretion Dynamics of UBB^+1^ Under Cycloheximide Treatment”. (**A and B**) WT HeLa cells expressing MYC-UBB⁺¹ were treated with 100 μg/mL cycloheximide (CHX) alone or in combination with Bafilomycin A1 (100 nM) or Velcade (100 nM) for the indicated time points (0, 4, and 8 h). Whole cell lysates (WCL) **(A)** and conditioned media **(B)** were collected and immunoblotted for UBB⁺¹. Equal volumes of conditioned media were harvested from comparable cell numbers across all conditions. Representative UBB⁺¹ blots are shown. Densitometric quantification of UBB⁺¹ levels from three independent experiments was performed. For WCL (A), UBB⁺¹ band intensities were normalized to GAPDH and expressed as percentage degradation relative to the 0 h time point (set as 100%). For conditioned media (B), UBB⁺¹ band intensities were quantified and plotted as secreted levels. Statistical comparisons were performed between CHX-alone and each of the inhibitor-treated conditions (CHX + Bafilomycin A1 or CHX + Velcade) at each time point. Comparisons at 0 h are indicated by round dotted lines, at 4 h by solid lines, and at 8 h by square dotted lines in the graph. Bar graphs represent mean ± SD from three biologically independent experiments. Statistical significance was determined using one-way ANOVA followed by Tukey’s multiple comparisons test. For all statistical tests: *, **, and ***, indicate p < 0.05, 0.01, and 0.001, respectively; ns = non-significant. **(C–F)** Conditioned media **(C, D)** and whole-cell lysates (WCL; **E, F**) were collected from wild-type (WT) and *sec22b* KO HeLa cells expressing MYC-UBB⁺¹ after treatment with 100 μg/mL cycloheximide (CHX) alone or in combination with Bafilomycin A1 (100 nM) for the indicated time points (0 h and 8 h). Equal volumes of conditioned media were harvested from comparable cell numbers across all conditions and immunoblotted for UBB⁺¹. For WCL (**E, F**), GAPDH was used as a loading control. UBB⁺¹ band intensities were normalized to GAPDH and expressed as a percentage of the 0 h time point (set to 100%). Representative immunoblots and densitometric quantification from three independent biological replicates are shown. For conditioned media (**C, D**), statistical comparisons were made between WT and *sec22b* KO cells for each of the following conditions: CHX 0 h (round dotted lines), CHX 8 h (solid lines), CHX + Bafilomycin A1 (8 h) (square dotted lines). Bar graphs represent, mean ± SD from three independent experiments. Statistical significance was assessed using an unpaired two-tailed t-test to compare WT and *sec22b* KO samples at each respective time point. Asterisks denote significance thresholds: *, **, and *** correspond to p < 0.05, p < 0.01, and p < 0.001, respectively; ns = not significant. In the graphs, WT and *sec22b* KO data are shown in black and gray, respectively.

### SQSTM1/p62 is required for efficient clearance of UBB+1 aggregates and its secretion

The uptake of ubiquitinated cargo into autophagosomes generally relies on specialized ubiquitin receptors. To test whether any of the known autophagy-related ubiquitin receptors are responsible for directing UBB^+1^ to secretion, we obtained individual and combinatorial knockouts of TAX1BP1, NBR1, OPTN, NDP52, and SQSTM1/p62 (**Fig. S3A-D**). A *penta* KO (5 KO) of all five receptors (*tax1bp1*/*optn*/*ndp52*/*nbr1*/*p62)* abolished secretion of UBB^+1^ (**Fig. 3A**). Since TAX1BP1 contains a UBZ (Ubiquitin-Binding Zinc Finger) domain with specificity for monoubiquitin, we considered it a potential candidate for association with the “single ubiquitin domain protein”, UBB^+1^. However, an individual *tax1bp1* KO showed no significant effect on UBB^+1^ secretion (**Fig. S3E**). Even a *triple* KO (TKO) of a subset of these receptors (*optn*/*ndp52*/*tax1bp1)* showed no significant effect on UBB^+1^ secretion (**Fig. 3B and 3C**). Surprisingly, the individual *p62* KO exhibited a complete loss of UBB^+1^ secretion (**Fig. 3D and 3E**), indicating that SQSTM1/p62 is essential and sufficient for UBB^+1^ secretion. In addition to its known role in aggrephagy (43-45), this study reveals that SQSTM1/p62 also plays a role in secretory autophagy.

**Figure 3.**
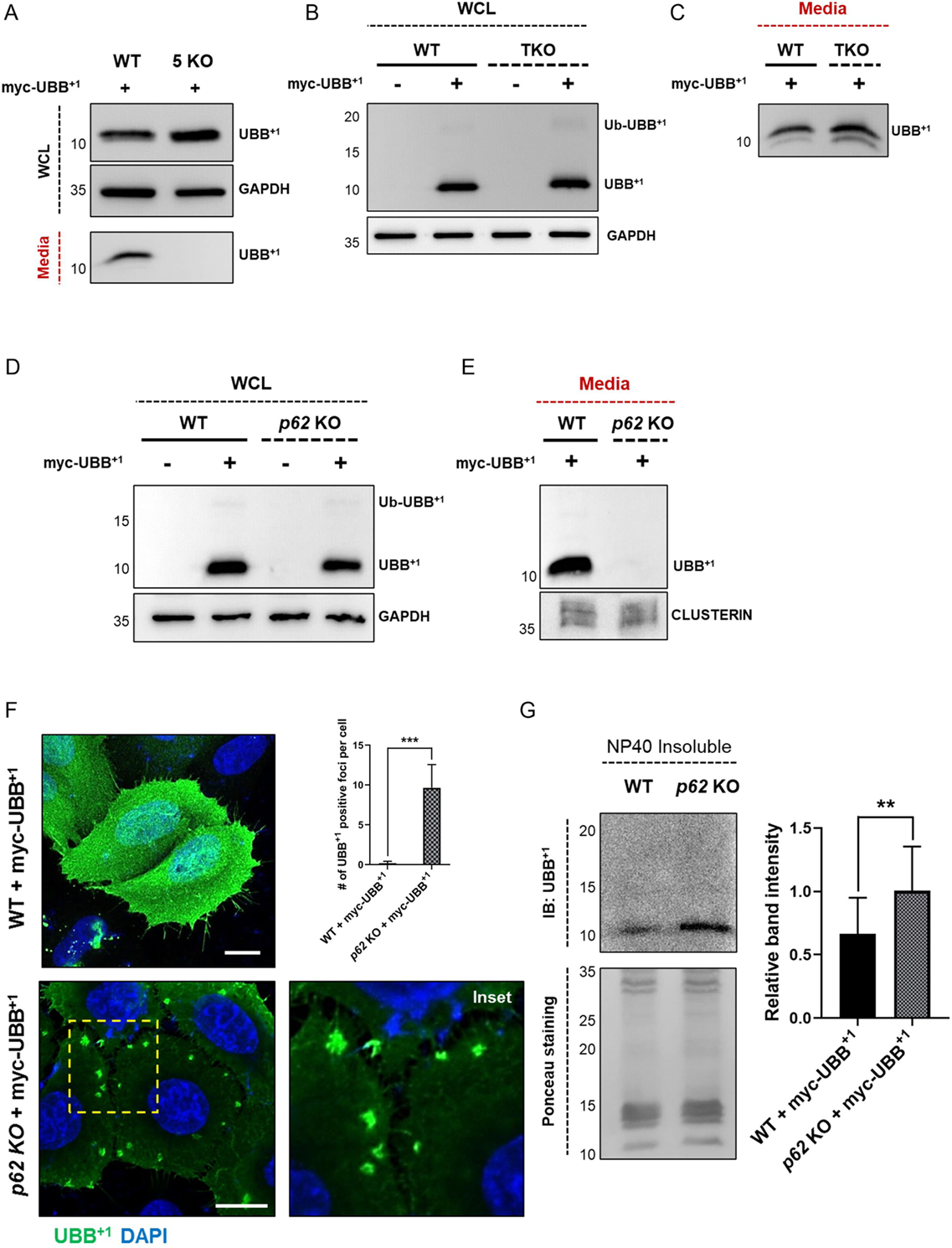
SQSTM1/p62 knockout hinders the efficient clearance of UBB^+1^. (**A-E**) Wild type (WT), *TAX1BP1/OPTN/NDP52/NBR1/p62* penta knockout (5KO), *OPTN/NDP52/TAX1BP1* triple knockout (TKO), and *p62* KO HeLa cells were transiently transfected with or without MYC-UBB⁺¹. After 24 h, cells were incubated with fresh DMEM for 16 h. Conditioned media and whole cell lysates (WCL) were resolved by 15% SDS-PAGE and immunoblotted for UBB⁺¹. GAPDH was used as the loading control for WCL. Conditioned media were collected in equal volumes from comparable cell numbers across all experimental conditions and immunoblotted for UBB⁺¹. **(F)** Representative immunofluorescence staining of WT and *p62* KO HeLa cells expressing MYC-UBB⁺¹. Cells were fixed, permeabilized, and immunostained with a monoclonal antibody against UBB⁺¹ (green), and nuclei were stained with DAPI (blue). Scale bars: 5 μm. Statistical significance was determined by unpaired two-tailed t-test, with data showing the mean ± SD from approximately 100 cells. **(G)** Representative immunoblot of UBB⁺¹ in insoluble fractions harvested from WT and *p62* KO HeLa cells transfected with MYC-UBB⁺¹ plasmid. Cells were lysed with NP40-lysis buffer, clarified at 15,000 x g and the pellet resuspended in 1% SDS. Ponceau staining was used for loading control. Data represents the mean ± SD from at least three independent experiments. For all statistical tests, *, **, ***, p < 0.05, 0.01, and 0.001, respectively.

Having uncovered an exclusive role for SQSTM1/p62 in UBB^+1^ secretion we turned our attention to the intracellular fate of UBB^+1^. How is its accumulation managed in the absence of secretion? Amorphous UBB^+1^ clusters were visualized in a KO of SQSTM1/p62 by immunofluorescence (**Fig. 3F**). Both *p62* KO and *penta* KO (**Fig. S3F**) cells accumulated UBB⁺¹-positive material, showing a similar spread-out distribution, suggesting that loss of SQSTM1/p62 alone is sufficient to disrupt UBB⁺¹. Does the involvement of SQSTM1/p62 in elimination of these UBB^+1^ puncta imply that intracellular UBB^+1^ is prone to aggregation? Indeed, the UBB^+1^ protein accumulated in insoluble fractions of *p62* KO cell extracts (**Fig. 3G**). To conclude, *SQSTM1*/*p62* deficient cells displayed a significant block in UBB^+1^ aggregate clearance, highlighting SQSTM1/p62’s role as a specific receptor for UBB^+1^, potentially for both degradative and secretory autophagy.

### Functional association between SQSTM1/p62 and the ubiquitin domain of UBB^+1^

Since SQSTM1/p62 interferes with UBB^+1^ aggregate formation on the one hand and is required for its secretion on the other hand, we explored the potential for direct interactions between these two proteins. As shown in **Figure 4A**, immunofluorescence of UBB^+1^-expressing HeLa cells revealed co-localization with endogenous SQSTM1/p62 in the form of intracellular puncta. However, a significant portion of UBB^+1^ was delocalized throughout the cell. This could suggest that the levels of endogenous SQSTM1/p62 were insufficient to associate with the heterologously expressed UBB^+1^. To address this, we repeated the experiment in cells overexpressing SQSTM1/p62. Immunofluorescence analysis of co-expressing cells showed prominent perinuclear puncta containing both proteins.

**Figure 4.**
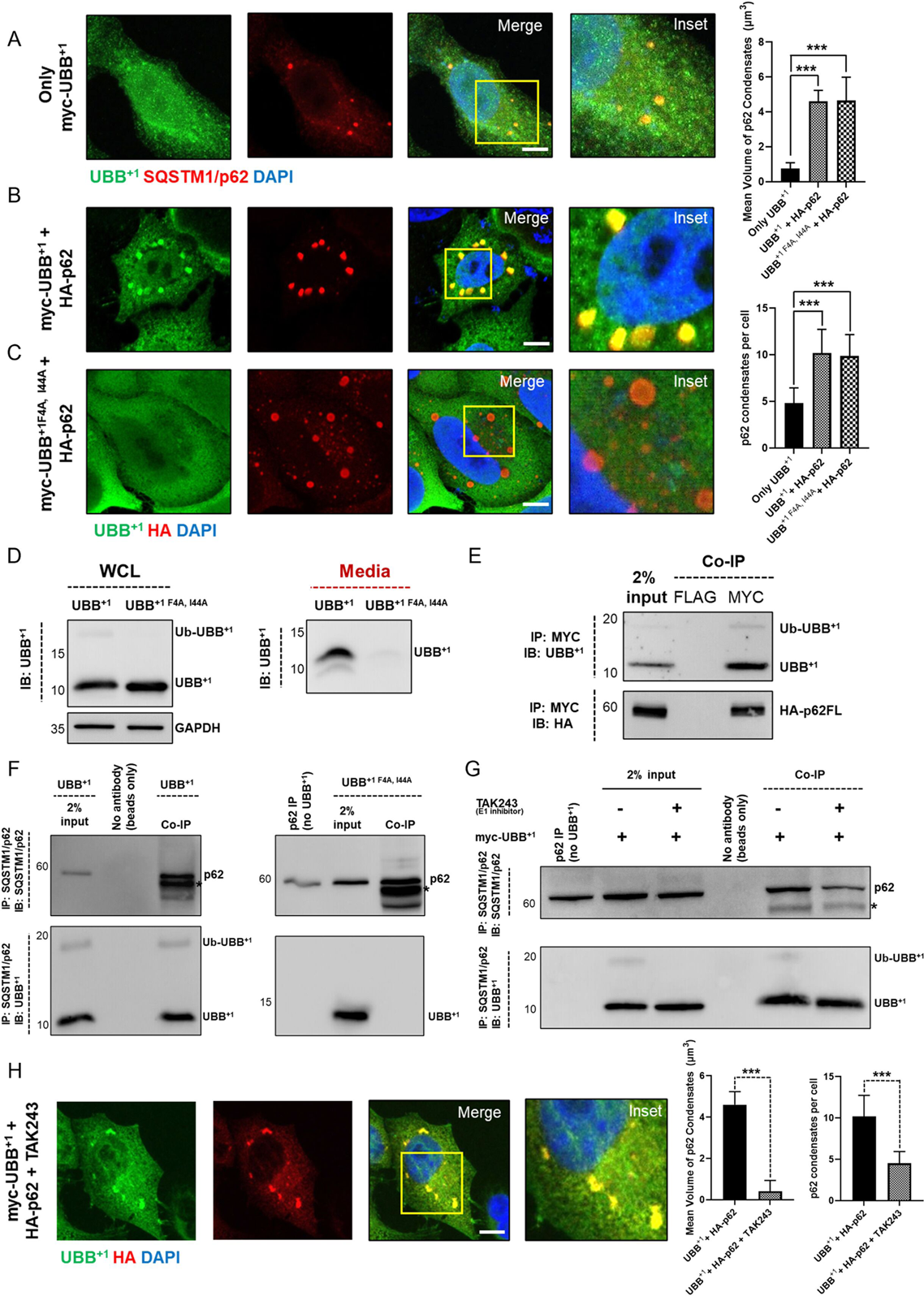
Evidence of direct interaction between SQSTM1/p62 and UBB^+1^. **(A)** Immunofluorescence staining of endogenous SQSTM1/p62 in HeLa cells expressing MYC-UBB⁺¹. Cells were grown in complete media, fixed, permeabilized, and stained with monoclonal antibodies for UBB⁺¹ (green) and SQSTM1/p62 (red). Merged images show colocalization, with nuclei stained using DAPI (blue). Scale bar: 2 μm. **(B)** Immunofluorescence staining of overexpressed HA-p62 in HeLa cells co-expressing MYC-UBB⁺¹. Cells were fixed, permeabilized, and stained with monoclonal antibodies for UBB⁺¹ (green) and HA (red). Merged images are shown, with nuclei counterstained using DAPI (blue). Scale bar: 2 μm. **(C)** Immunofluorescence staining of HeLa cells expressing HA-p62 and MYC-UBB⁺¹ ^F4A, I44A^. Superscript denotes point mutations (F4A and I44A) introduced into the UBB⁺¹ protein. UBB⁺¹ ^F4A, I44A^ showed no clear colocalization with HA-p62 condensates. Scale bar: 2 μm. Quantification of condensate volume per cell and condensate number per cell was performed using Imaris-based image analysis. Data represents a minimum of 100 cells per condition, pooled from three independent experiments. In both the condensate volume and condensate number graphs, statistical comparisons were made between the UBB⁺¹-only condition and each of the following: UBB⁺¹ + HA-p62 and UBB⁺¹ ^F4A, I44A^ + HA-p62, as indicated by solid lines. Bar graphs represent mean ± SD. Statistical significance was determined using one-way ANOVA followed by Tukey’s multiple comparisons test. For all statistical tests: *, **, and ***, indicate p < 0.05, 0.01, and 0.001, respectively. **(D)** Immunoblot analysis of conditioned media and whole cell lysates (WCL) from HeLa cells expressing MYC-UBB⁺¹ or the hydrophobic patch mutant MYC-UBB⁺¹ ^F4A, I44A^. Cells were transfected for 24 h and then incubated in fresh DMEM for 16 h. Conditioned media and WCL were resolved by 15% SDS-PAGE and immunoblotted using specific antibodies against UBB⁺¹. GAPDH was used as a loading control for WCL. Conditioned media were collected in equal volumes from comparable cell numbers across all experimental conditions to ensure consistent comparison of secreted protein levels. **(E)** Co-immunoprecipitation (co-IP) of HA-tagged p62 with MYC-UBB⁺¹ in HeLa cells. Cells were co-transfected with HA-p62 and MYC-UBB⁺¹, and lysates were subjected to immunoprecipitation under non-denaturing conditions using MYC- or FLAG-conjugated magnetic beads. Bound proteins were eluted and analysed by immunoblotting (IB) with anti-HA and anti-UBB⁺¹ antibodies to assess interaction between p62 and UBB⁺¹. **(F)** Co-immunoprecipitation of endogenous SQSTM1/p62 with MYC-UBB⁺¹ in HeLa cells. Lysates from cells expressing MYC-UBB⁺¹ were immunoprecipitated using an anti-p62 antibody under non-denaturing conditions. Immunoprecipitates were resolved by SDS-PAGE and probed with anti-p62 and anti-UBB⁺¹ antibodies. The anti-UBB⁺¹ antibody detected both ubiquitinated and non-ubiquitinated forms of UBB⁺¹, while the immunoglobulin heavy chain is visible as a ∼50 kDa band (indicated by an asterisk). Negative controls included: (1) a beads-only (no antibody) IP control to evaluate non-specific binding, and (2) IP from untransfected cells lacking MYC-UBB⁺¹ expression to assess background signal. **(G)** Co-immunoprecipitation of endogenous SQSTM1/p62 with MYC-UBB⁺¹ following ubiquitination inhibition by TAK243. HeLa cells were transfected with MYC-UBB⁺¹ and treated with TAK243 (1 µM, 3 h) to inhibit ubiquitin-activating enzyme (E1) activity. Lysates were immunoprecipitated using an anti-p62 antibody and analysed by immunoblotting with anti-p62 and anti-UBB⁺¹ antibodies. Both ubiquitinated and non-ubiquitinated forms of UBB⁺¹ were detected in the input; however, only the non-ubiquitinated form co-precipitated with p62 following TAK243 treatment. The antibody heavy chain (∼50 kDa) is marked with an asterisk. Negative controls included beads-only IP and IP from untransfected cells to rule out non-specific interactions. **(H)** Immunofluorescence staining of HeLa cells co-expressing HA-p62 and MYC-UBB⁺¹ after 3 h of TAK243 treatment (same as in panel G). Cells were fixed, permeabilized, and stained with monoclonal antibodies for UBB⁺¹ (green) and HA (red). Nuclei were counterstained using DAPI (blue). Merged images and zoomed insets show the intracellular distribution and colocalization of p62 and UBB⁺¹ following ubiquitination blockade. Scale bar: 2 μm. Quantification of p62 condensate volume and condensate number per cell was performed from the same experimental set shown in panel (H), using Imaris image analysis software. Data was obtained from a minimum of 100 cells per condition, pooled from three independent experiments. Statistical comparisons were made between the UBB⁺¹ + HA-p62 WT and UBB⁺¹ + HA-p62 WT + TAK243 conditions. In the graphs, comparisons are indicated by square-dotted lines. Bar graphs represent mean ± SD. Statistical significance was assessed using one-way ANOVA followed by Tukey’s multiple comparisons test. For all statistical tests: *, **, and ***, indicate p < 0.05, 0.01, and 0.001, respectively.

Previous studies have reported the formation of membrane-less inclusion bodies composed of SQSTM1/p62 and ubiquitinated substrates upon ectopic expression of SQSTM1/p62, termed ‘p62-bodies’ or ‘sequestosomes’ (45, 46). We similarly observed the formation of such p62-bodies when SQSTM1/p62 was overexpressed in the current study (**Fig. S4A**). However, in UBB^+1^-expressing cells, these bodies were organized in an ordered, perinuclear pattern (**Fig. 4B and S4B**) strengthening our conclusion that the two proteins are co-localized. The presence of a potential (ubiquitinated) substrate, such as UBB^+1^, appears to influence the intracellular localization and organization of these p62-bodies.

SQSTM1/p62 is a critical ubiquitin-binding protein that facilitates the shuttling of ubiquitinated cargo towards autophagosomes. To elucidate the molecular mechanisms of recognition, we mutated key recognition elements within the ubiquitin domain of UBB^+1^. Previous studies have identified two hydrophobic surface residues on ubiquitin, Phe4 (F4) and Ile44 (I44), that are recognized directly by the overwhelming majority of Ub receptors (47). To investigate the role of these hydrophobic patches in UBB^+1^ recognition, we generated a double substitution F4A and I44A in UBB^+1^ and co-expressed UBB^+1^ ^F4A, I44A^ with SQSTM1/p62 in HeLa cells. Immunofluorescence showed a complete loss in co-localization between SQSTM1/p62 and UBB^+1^ ^F4A, I44A^ and disruption of the characteristic perinuclear pattern of UBB^+1^-containing *p62-bodies* (**Fig. 4C**). A significant reduction in secretion of the hydrophobic patch mutant compared to WT UBB^+1^ (**Fig. 4D**) further supported the importance of the ubiquitin domain for recognition of UBB^+1^ by SQSTM1/p62 and for its secretion. Additionally, the hydrophobic patch mutant led to undetectable ubiquitination on UBB^+1^ ^F4A, I44A^ (**Fig. 4D**). Therefore, our next question was whether the binding of SQSTM1/p62 is directly to the surface of UBB^+1^ or to its ubiquitinated form.

Co-immunoprecipitation (co-IP) was employed to investigate whether SQSTM1/p62 preferentially binds to the ubiquitinated or non-ubiquitinated forms of UBB^+1^. First, immunoprecipitation (IP) of MYC-tagged UBB⁺¹ co-precipitated HA-p62 (**Fig. 4E**), indicating a specific association between the two proteins. Next, precipitation of endogenous SQSTM1/p62 co-immunoprecipitated UBB⁺¹ (**Fig. 4F**), further supporting a physical association. Consistent with a requirement for an intact ubiquitin surface, the hydrophobic-patch mutant MYC-UBB⁺¹ ^F4A, I44A^ failed to co-precipitate endogenous SQSTM1/p62 and showed negligible colocalization by immunofluorescence (**Fig. 4F**).

To test whether additional ubiquitin chains are required for this association, we blocked all cellular ubiquitination by inhibiting the ubiquitin-activating enzyme (E1). This treatment depleted polyubiquitin conjugates and eliminated ubiquitination of UBB⁺¹ (**Fig. S4C and S4D**). The remaining unmodified UBB⁺¹ nevertheless interacted with SQSTM1/p62 in co-IP assays and colocalized in cells (**Fig. 4G and 4H**). Across all co-IP’s, we noticed that the ratio between ubiquitinated and unmodified UBB^+1^ in the co-IP eluate was unaltered relative to the ratio in whole cell extract (**Fig. 4F**) indicating that SQSTM1/p62 does not have a marked preference for one form or the other. Collectively, these findings identify SQSTM1/p62 as an interacting partner of UBB^+1^ independent of its ubiquitination state. Through this association, SQSTM1/p62 plays a crucial role in regulating intracellular levels of UBB^+1^ by alleviating its aggregation and promoting its secretion.

### Mapping SQSTM1/p62 domains involved in UBB^+1^ binding and secretion

As a scaffold protein, SQSTM1/p62 has multiple functional domains, through which it associates with its interacting partners. Beyond a C-terminal UBA domain that binds ubiquitinated cargo (15, 48-50), SQSTM1/p62 contains an N-terminal PB1 domain which drives self-oligomerization, and an LC3-interacting region (LIR) motif mediating the interaction with ATG8 family proteins (51-53). To better understand the role of these domains in regulating UBB^+1^ secretion, we constructed SQSTM1/p62 lacking its PB1 domain (p62ΔPB1), a version with a non-functional LIR motif (p62 ^W340A, L343A^), and a UBA domain mutant (p62 ^M404V^), which disrupts hydrophobic interactions essential for ubiquitin binding. All constructs were HA-tagged and expressed in *p62* KO HeLa cells co-expressing UBB⁺¹ (**Fig. 5A)**.

**Figure 5.**
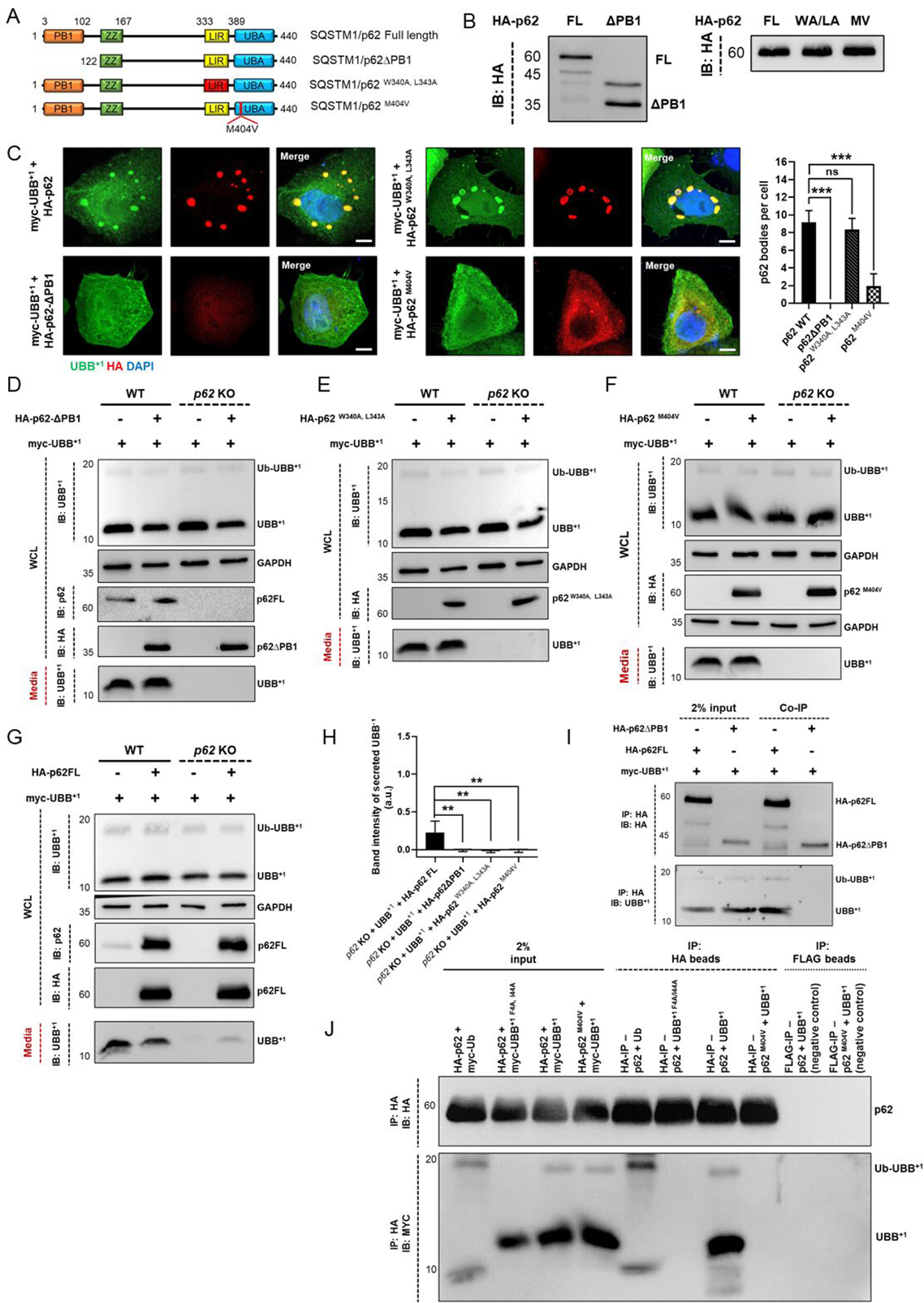
SQSTM1/p62 domains required for functional association with UBB^+1.^ **(A)** Schematic representation of functional domains in full-length SQSTM1/p62 (FL), the ΔPB1 deletion mutant (lacking the N-terminal PB1 oligomerization domain; p62ΔPB1), the LC3-interacting region (LIR) mutant (W340A, L343A; denoted p62 ^W340A, L343A^), and the UBA domain point mutant (p62 ^M404V^). **(B)** HeLa cells were transiently transfected with plasmids encoding C-terminally HA-tagged SQSTM1/p62 constructs: FL, ΔPB1, WA/LA, and MV. After 24 h, whole-cell lysates were prepared and subjected to SDS-PAGE followed by immunoblotting using an anti-HA antibody to assess expression levels. All constructs were expressed at comparable levels, confirming that functional differences observed in secretion and localization assays (Fig. 5) are not due to unequal expression. The predicted molecular weight of full-length p62 is 47 kDa; however, it migrates above 60 kDa due to post-translational modifications. The truncated ΔPB1 construct has a calculated MW of 35.5 kDa, also appearing as a modified higher MW form. **(C)** Immunofluorescence staining of HeLa cells co-expressing MYC-UBB⁺¹ with either full-length HA-p62 (FL), HA-p62ΔPB1 (lacking the N-terminal PB1 oligomerization domain), HA-p62 ^W340A, L343A^ (LC3-interacting region mutant; denoted WA/LA), or HA-p62 ^M404V^ (UBA domain mutant defective in ubiquitin binding). Cells were fixed 24 h post-transfection, permeabilized, and stained with monoclonal antibodies specific for HA (red) and UBB⁺¹ (green); nuclei were counterstained with DAPI (blue). The ΔPB1 mutant failed to form *p62-bodies* due to impaired oligomerization, while the M404V mutant showed markedly reduced colocalization and condensate formation with UBB⁺¹ due to loss of UBA-mediated binding. In contrast, both FL and WA/LA constructs retained the ability to form p62-positive condensates. Scale bar: 2 μm. **Right:** Quantification of p62-positive condensates per cell under each condition. Data represent mean ± SD from >100 cells per group pooled from three biologically independent experiments. Statistical significance was determined using one-way ANOVA followed by Tukey’s multiple comparisons test. For all statistical tests: *p < 0.05, **p < 0.01, ***p < 0.001. **(D–G)** Immunoblot analysis of conditioned media and whole-cell lysates (WCL) from WT and *p62* KO HeLa cells co-transfected with MYC-UBB⁺¹ and either HA-p62ΔPB1 **(D)**, HA-p62 *W340A, L343A* **(E)**, HA-p62 *M404V* **(F)**, or full-length HA-p62 (FL) **(G)**. After 24 h of transfection, cells were incubated with fresh DMEM for an additional 16 h. Conditioned media and WCL were collected, resolved by 10% or 15% SDS-PAGE, and immunoblotted with the indicated antibodies. GAPDH was used as a loading control for WCL. Equal volumes of conditioned media were harvested from comparable cell numbers across all experimental conditions. Representative immunoblots showing UBB⁺¹ levels in the media are displayed. **(H)** Densitometric quantification of UBB⁺¹ levels in the conditioned media across p62 KO cells co-expressing MYC-UBB⁺¹ with either full-length HA-p62 or the indicated p62 mutant constructs (ΔPB1, W340A/L343A, or M404V). Quantification is based on three biologically independent experiments. Statistical comparisons were performed between the full-length p62 condition and each of the mutant constructs. Bar graphs represent mean ± SD. Statistical significance was assessed using one-way ANOVA followed by Tukey’s multiple comparisons test. For all statistical tests: *p < 0.05, **p < 0.01, ***p < 0.001; ns = non-significant. **(I)** Co-immunoprecipitation (Co-IP) of HA-tagged SQSTM1/p62 constructs with MYC-UBB⁺¹ from HeLa cell lysates. Cells were co-transfected with MYC-UBB⁺¹ and either full-length HA-p62 FL or HA-p62ΔPB1. After 24 h, cells were lysed under non-denaturing conditions, and immunoprecipitation was performed using anti-HA magnetic beads. Elution was carried out using HA peptide. Immunoprecipitates were analyzed by SDS-PAGE and immunoblotting (IB) with anti-HA and anti-UBB⁺¹ antibodies. **(J)** Co-immunoprecipitation of MYC-UBB⁺¹ or MYC-UBB⁺¹ mutants with HA-p62 or HA-p62M404V from HeLa cell lysates. Lanes 1–4 show 2% input: lane 1, MYC-UB and HA-p62 (positive control); lane 2, MYC-UBB⁺¹ ^F4A, I44A^ mutant and HA-p62 (negative control); lane 3, MYC-UBB⁺¹ and HA-p62; lane 4, MYC-UBB⁺¹ and HA-p62 ^M404V^. Lanes 5–8 display HA immunoprecipitations using anti-HA magnetic beads: lanes 5–7 with HA-p62; lane 8 with HA-p62 ^M404V^. Lanes 9–10 serve as additional negative controls in which lysates were incubated with anti-FLAG magnetic beads: lane 9, HA-p62 + MYC-UBB⁺¹; lane 10, HA-p62M404V + MYC-UBB⁺¹. Immunoprecipitates were probed using anti-MYC and anti-HA antibodies. The results confirm a specific interaction between wild type p62 and UBB⁺¹, which is abolished when either the hydrophobic patch of UBB⁺¹ (F4A, I44A) or the Met404 residue in the UBA domain of p62 is mutated.

In contrast to full-length (FL) SQSTM1/p62, overexpression of truncated SQSTM1/p62 (p62ΔPB1) lacking the self-oligomerization property did not form *p62-bodies* with UBB^+1^ (**Fig. 5B**), emphasizing the essential role of the PB1 domain in body formation. By contrast, the construct lacking a functional LIR motif (p62 ^W340A, L343A^) displayed the characteristic perinuclear *p62-bodies* (**Fig. 5B**), supporting a model whereby *p62-bodies* form upstream to autophagosomes. Strikingly, the M404V mutation in the UBA domain abolished p62’s ability to co-localize with UBB⁺¹, indicating that UBA-dependent binding to ubiquitinated UBB⁺¹ is critical for cargo engagement (**Fig. 5B**).

In *p62* KO cells, expression of UBB⁺¹ alone resulted in a complete loss of secretion compared to WT cells. Overexpression of HA-p62FL in the KO background restored secretion, demonstrating that full-length p62 is sufficient to rescue this defect (**Fig. 5G**). In contrast, none of the mutant constructs (p62ΔPB1, p62W340A/L343A, or p62M404V), expressed at comparable levels (**Fig. 5B**), were able to rescue secretion (**Fig. 5D–F**). These findings underscore the requirement for all three functional modules (PB1, LIR, and UBA) in facilitating efficient UBB⁺¹ sequestration and routing into the secretory autophagy pathway. To exclude the possibility that these results reflect artifacts of SQSTM1/p62 overexpression, we performed dose-dependent titration experiments in both WT and *p62* KO cells (**Fig. S5A–C**). Increasing plasmid concentrations of HA-p62FL (0 ng, 100 ng, 250 ng, 500 ng, 1 µg) demonstrated that at moderate expression levels, HA-p62FL reproducibly enhanced UBB⁺¹ secretion, whereas very high overexpression led to a dominant-negative effect through the formation of large p62-bodies (**Fig. S5B**). Expression levels of the different HA-p62 constructs were validated by both immunofluorescence and western blotting, confirming comparability across constructs (**Fig. 5B**). Together, these results establish HA-p62FL as a robust rescue reference and demonstrate that disruption of the PB1, LIR, or UBA domains abolishes the ability of SQSTM1/p62 to restore UBB⁺¹ secretion.

Without a functional LIR motif, SQSTM1/p62 is not expected to shuttle ubiquitinated cargo to LC3-containing phagophores, explaining the inability of this mutant to support UBB^+1^ secretion (**Fig. 5E**). However, why would a SQSTM1/p62 construct lacking a self-oligomerization (PB1) domain be unable to promote the secretion of UBB^+1^ (**Fig. 5D**)? In light of the inability of p62ΔPB1 to co-localize with UBB^+1^ (**Fig. 5C**), we proposed that the PB1 may potentially participate in UBB^+1^ binding. To further define the binding requirements, we next evaluated the role of the UBA domain using an HA-p62^M404V^ mutant known to abolish binding to polyubiquitin (54-56). Co-immunoprecipitation and immunofluorescence assays showed that HA-p62^M404V^ failed to interact with or form p62 bodies with UBB⁺¹ (**Fig. 5C and 5J**). Moreover, when expressed in p62 KO cells, this mutant was unable to rescue UBB⁺¹ secretion (**Fig. 5F**), in contrast to HA-p62 FL (**Fig. 5G**). Importantly, Western blot analysis demonstrated that expression levels of HA-p62 FL, HA-p62ΔPB1, HA-p62 ^W340A, L343A^, and HA-p62 ^M404V^ were all comparable and within moderate ranges, suggesting that the failure to rescue secretion was specifically due to impaired function rather than artifacts from excessive overexpression (**Fig. 5B**). These results strongly support the notion that the intact UBA domain and specifically residue M404 is essential for p62-mediated UBB⁺¹ recognition, aggregation, and secretion.

It has been reported that the deletion of the PB1 domain or oligomerization-inhibiting mutations resulted in reduced interaction with both LC3B and ubiquitin in pull-down assays, suggesting that oligomerization may increase the interaction with these binding partners (8, 45). Therefore, we tested whether p62ΔPB1 could bind to UBB^+1^. A pulldown assay of HA-p62ΔPB1 showed no evidence of interaction with UBB^+1^ (**Fig. 5G**). Apparently, due to the loss of oligomerization of SQSTM1/p62, p62ΔPB1 was insufficient to trap UBB^+1^ despite containing an intact UBA domain. The inherent property of SQSTM1/p62 to oligomerize and form tandem interactions with multiple Ub domains may underline the body formation we witnessed with UBB^+1^. Lacking the ability to self-oligomerize, p62ΔPB1 can only bind a single unit at a time, which for a single ubiquitin domain protein such as UBB^+1^ is insufficient, apparently, for tight binding. For this reason, we propose that p62ΔPB1 was unable to sequester UBB^+1^ or package it into autophagosomes.

To gain additional structural insight into the interaction between UBB⁺¹ and the UBA domain of SQSTM1/p62, we used AlphaFold3 to model a complex between the two proteins based on their primary sequences (**Fig. S5D and S5E**). The predicted structure revealed that residue M404 in the p62 UBA domain is positioned near to a conserved hydrophobic patch on UBB⁺¹ comprising residues Isoleucine 44, Leucine 8, and Valine 70. When this methionine was mutated to valine (V404), the predicted distance between V404 and these hydrophobic residues increased, suggesting a loss of stabilizing hydrophobic contacts. These structural observations are consistent with our biochemical and imaging data, which show that the M404V mutation disrupts UBB⁺¹ binding, impairs p62-body formation, and blocks UBB⁺¹ secretion (**Fig. 5C and 5F**). This supports a critical role for M404 in mediating UBB⁺¹ recognition by p62 through hydrophobic interaction.

## Discussion

Accumulation of UBB^+1^ is toxic to cells, particularly neurons, where it disrupts cellular homeostasis. Recently, UBB⁺¹ has been identified as a key factor in inducing AD-like pathology (25). Cells appear to have evolved multiple mechanisms to counteract UBB⁺¹-associated stress by reducing its intracellular levels. UBB⁺¹ can be partially degraded by the proteasome (57, 58), and findings from this study suggest that lysosomes also contribute to its intracellular turnover. Cells can also actively secrete UBB^+1^ via an unconventional autophagosome-like vesicle-mediated pathway (31).

Overexpression of UBB⁺¹ in cultured cells enables detailed mechanistic analysis of its trafficking and clearance, a widely used approach for studying aggregation-prone proteins implicated in neurodegeneration (31, 59, 60). Insights gained from such models have consistently mirrored findings in neurons and patient-derived tissues, thereby reinforcing the physiological relevance of our observations. In this study, we identified a novel function of secretory autophagy in mitigating UBB⁺¹ toxicity. Our findings show that UBB⁺¹ is actively regulated by an autophagy-mediated vesicle release mechanism, with the autophagy receptor SQSTM1/p62 playing a central role. SQSTM1/p62 promotes the sequestration of UBB⁺¹ into specialized *p62-bodies*, a process that occurs independently of UBB⁺¹’s ubiquitination status (**Fig. 3 and 4**). The N-terminal PB1 domain of SQSTM1/p62 plays a critical role in its oligomerization, allowing the formation of higher-order structures that are essential for the efficient sequestration of ubiquitinated cargo (45, 61). This oligomerization facilitates the clustering of ubiquitinated proteins into *p62-bodies*, which act as scaffolds for selective autophagy. Likewise, our results revealed that the PB1 domain is essential for the sequestration of UBB⁺¹ by SQSTM1/p62, and in its absence, *p62-body* formation was disrupted (**Fig. 5**). The measured affinity of the UBA domain of SQSTM1/p62 for monoubiquitin is relatively weak (*K_d_ = 540 µM*) (15). The PB1 domain facilitates higher ordered oligomers of SQSTM1/p62, which interact with multiple ubiquitin molecules (45), which in our case explains how SQSTM1/p62 can bind the non-ubiquitinated form of UBB^+1^.

A limitation of these assays is that they rely on overexpression of p62 constructs. While our titration experiments confirmed that moderate expression levels reproducibly restore UBB⁺¹ secretion, we also observed that very high overexpression can reduce secretion due to dominant-negative effects caused by excessive *p62-body* formation. These findings indicate that dosage is an important variable when interpreting rescue experiments and should be considered in future studies using more physiological expression systems.

The ensuing *p62-bodies* help isolate UBB⁺¹ from the cytosol, preventing its aggregation and facilitating its recruitment into autophagosome-like vesicles for secretion (**Fig. 6)**. SQSTM1/p62 is shared between degradative and secretory autophagy. Recent studies have begun to uncover roles for SQSTM1/p62 in the secretion of other neurodegenerative disease–related proteins. For example, a recent report (62) demonstrated that SQSTM1/p62 facilitates the unconventional secretion of alpha-synuclein from neurons using mechanisms that overlap with those identified here for UBB⁺¹. In that study, SQSTM1/p62 was shown to interact with intracellular alpha-synuclein and direct it to autophagosome-derived secretory vesicles, independent of canonical lysosomal degradation. This emerging role of SQSTM1/p62 as a gatekeeper not only for degradation but also for secretion of misfolded or aggregation-prone proteins underscores a broader neuroprotective strategy employed by cells. Our findings extend this concept to UBB⁺¹, an aberrant ubiquitin variant associated with AD, highlighting that SQSTM1/p62-driven secretory autophagy may be generalizable for mitigating proteotoxic stress across multiple neurodegenerative contexts. These parallels suggest that similar vesicular routes may govern the disposal of different pathological proteins, offering potential targets for modulating disease progression.

**Figure 6.**
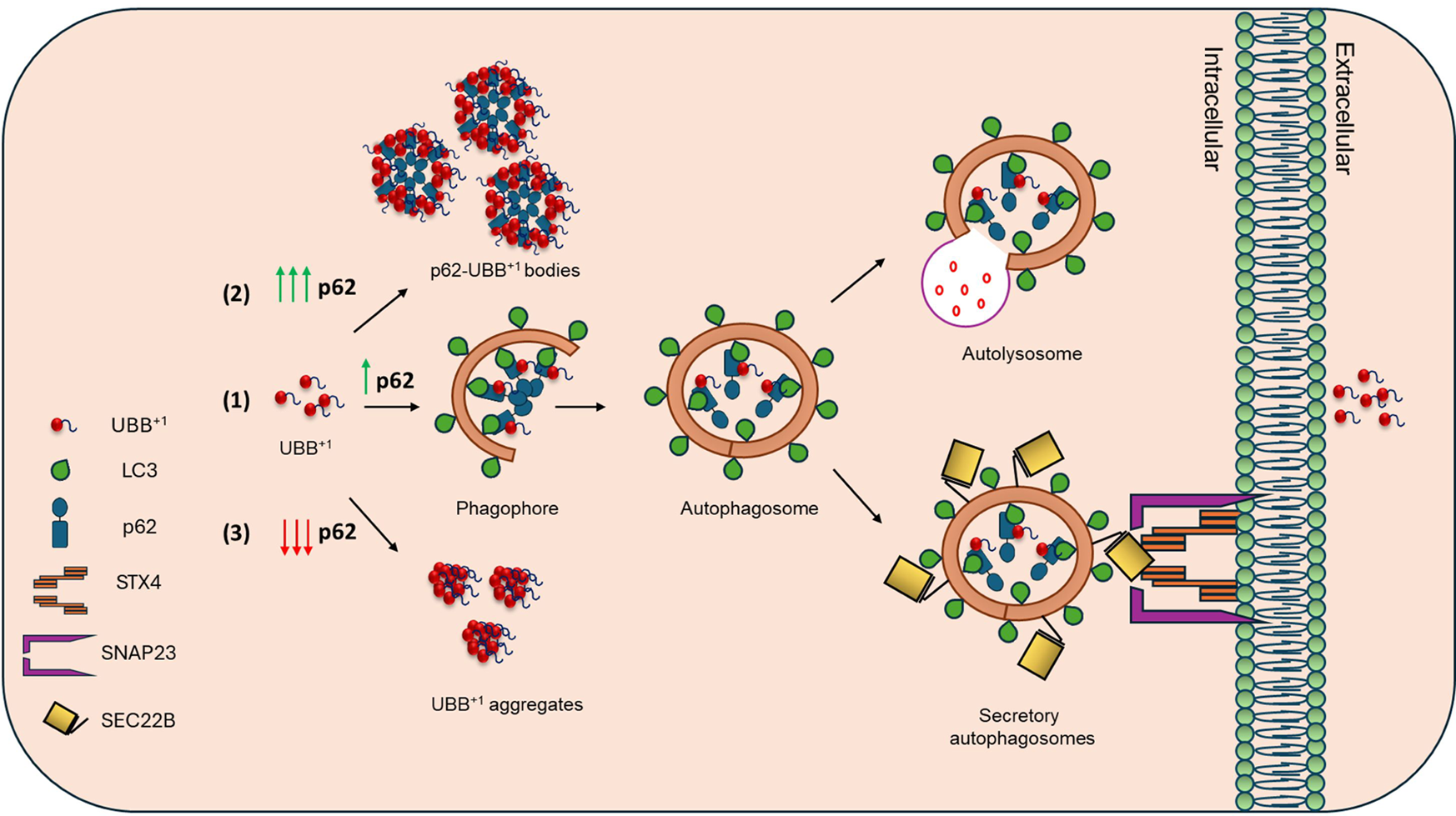
SQSTM1/p62 and the Cellular fates of UBB^+1^. SQSTM1/p62 critically affects both the intra- and extra-cellular fates of UBB^+1^: (1) Autophagic clearance facilitated by SQSTM1/p62 and LC3, leading to either lysosomal degradation or extracellular expulsion via autophagosome-plasma membrane fusion mediated by SEC22B and STX4/SNAP23. (2) Sequestration into p62-induced round bodies when SQSTM1/p62 is elevated relative to levels of LC3. (3) Aggregation and intracellular retention in the absence – or insufficient levels – of SQSTM1/p62.

The mere presence of UBB⁺¹ increases autophagic vesicle formation (31, 63). Supporting this observation, our results show that intracellular UBB⁺¹ accumulation in the absence of SQSTM1/p62 strongly induces LC3B-II conversion, indicating that aggregated UBB⁺¹ itself is sufficient to trigger autophagic responses independently of SQSTM1/p62 (**Fig. S3G**). Future studies should further explore whether such UBB⁺¹- induced autophagy is predominantly degradative or can also stimulate SEC22B-mediated secretory pathways under different cellular conditions, such as nutrient deprivation. Once packaged into autophagic vesicles aided by SQSTM1/p62, a subset of UBB⁺¹-containing autophagosomes bypass lysosomal degradation.

Depletion of SEC22B, an R-SNARE involved in unconventional protein secretion [3, 31], also reduced UBB⁺¹ secretion. In the absence of SEC22B, UBB⁺¹ colocalized with LC3-containing autophagosomes (**Fig. 1**). Although SEC22B plays a significant role in UBB⁺¹ secretion, our findings suggest that additional compensatory secretory pathways may exist, particularly under conditions where intracellular degradation is impaired. This may either involve other SNAREs or non-canonical vesicular trafficking pathways independent of classical secretory autophagy.

Knockdown of syntaxin 17 (STX17), a Qa-SNARE essential for autophagosome-lysosome fusion [35], did not impair UBB⁺¹ secretion (**Fig. S1C**). Instead, we identified syntaxin 4 (STX4) as the critical Qa-SNARE mediating the fusion of UBB⁺¹-containing vesicles with the plasma membrane. Knockdown of STX4 significantly reduced UBB⁺¹ secretion, whereas knockdown of syntaxin 3 (STX3) had no discernible effect. In addition, we explored the involvement of SNAP23 and SNAP29, known regulators of vesicle-plasma membrane fusion. Knockdown experiments revealed that depletion of SNAP23 significantly impaired UBB⁺¹ secretion, like the effect observed with STX4 knockdown, whereas SNAP29 depletion showed no significant effect (**Fig. 1D**). While some autophagosomes target UBB⁺¹ for lysosomal degradation, others are secreted through an unconventional autophagy-related pathway mediated by a SNARE complex including SEC22B, STX4, and SNAP23, facilitating the fusion of UBB⁺¹-loaded vesicles with the plasma membrane (**Fig. 6**).

This work identifies UBB¹ as a novel cargo for secretory autophagy specifically targeted through its ubiquitin domain. UBB⁺¹ contains an internal, non-cleavable ubiquitin domain that is integral to its structure. In this study, we provide initial evidence suggesting that this ubiquitin domain participates in the secretion of UBB⁺¹ and may contribute to its recognition for secretory autophagy. Specifically, we demonstrate that UBB⁺¹ colocalizes with the autophagy receptor SQSTM1/p62 and that SQSTM1/p62 physically associates with UBB⁺¹ through its ubiquitin domain. Furthermore, disruption of the hydrophobic patch within this ubiquitin domain significantly impairs SQSTM1/p62 association and inhibits UBB⁺¹ secretion. Together, these findings support the idea that the internal ubiquitin domain of UBB⁺¹ plays an important role in its selective recruitment into the secretory autophagy pathway.

In summary, our study exposes a dual role of autophagy in both intracellular protein quality control and a distinct pathway for UBB⁺¹ secretion mediated by SQSTM1/p62 and SEC22B. Secretion appears to represent an active strategy for alleviating UBB⁺¹- induced cellular stress. While ubiquitin is traditionally viewed as a degradation signal, this work highlights a unique example of a ubiquitin-related protein driving secretory autophagy. This process represents a specialized form of secretory autophagy that requires a ubiquitin receptor, yet bypasses lysosomal degradation, utilizing the SEC22B-STX4/SNAP23 SNARE complexes for vesicle-plasma membrane fusion. These findings provide new insights into how cells manage UBB⁺¹ toxicity and highlight the broader implications of secretory autophagy in proteostasis, intercellular communication, and neurodegenerative disease progression. Future studies could focus on identifying additional factors regulating this pathway and exploring the extracellular ramifications of UBB⁺¹ in health and disease.

Secretion raises intriguing questions about the fate of UBB⁺¹ in the extracellular environment. While this mechanism may initially benefit the cell by alleviating intracellular stress, the extracellular propagation of UBB⁺¹ could have pathological consequences. The potential involvement of UBB⁺¹ in extracellular amyloid plaque formation or spreading of toxic effects warrants further investigation.

## Material and methods

### Chemicals and Antibodies

DL-dithiothreitol (DTT, D0632), iodoacetamide (I1149), IGEPAL CA-630 (NP40, I3021), bovine serum albumin (BSA, A7906), sodium dodecyl sulfate (SDS, L3771), Tween-20 (P9416), ethylenediaminetetraacetic acid disodium salt dehydrate (EDTA, E4884), bafilomycin A1 (B1793) and Velcade (504314) were purchased from Sigma-Aldrich. TAK-243 (MLN7243) (S8341), Venetoclax (ABT-199; HY-15531) was purchased from Selleckchem. Complete protease inhibitor cocktail (04693116001) was purchased from Roche Diagnostic.

Antibodies and their manufacturers were: rabbit anti-UBB^+1^ (WB 1:1000, IF 1:250 custom made by Sigma for our lab) (25, 31). Rabbit anti-GAPDH (WB 1:10,000, G9545), mouse anti-HA tag (WB 1:2000, IF 1:250, H3663) were from Sigma-Aldrich. Mouse anti-CLUSTERIN (WB 1:1000; 166907), mouse anti-UBIQUITIN (WB 1:1000, 8017), mouse anti-ATG5 (WB 1:1000, 133158), mouse anti-SEC22B (WB 1:1000, 101267), mouse anti-ACTIN (WB 1:1000, 47778) and mouse anti-TSG101 (WB 1:1000, 7964), mouse anti-SNAP23 (WB 1:1000, 166244), mouse anti-SNAP29 (WB 1:1000, 390602) were from Santa Cruz Biotechnology. Rabbit anti-LC3 (WB: 1:2000, 51520) was from Abcam. Mouse anti-SQSTM1/p62 (WB 1:1000, IF, 1:250 88588), mouse anti-STX4 (WB 1:1000, E6W7B), mouse anti-STX17 (WB 1:1000, D3D7H), rabbit anti-cleaved caspase 3 (WB 1:1000, 9661) were from Cell Signaling Technology. Rabbit anti-STX3 (WB 1:1000, Synaptic systems, 110032). Alexa Fluor 488- and 594- conjugated secondary antibodies were from Thermo Fisher (A21206 and A11032 respectively). Peroxidase-conjugated anti-mouse (WB 1:10,1000, 115-005-003 and anti-rabbit secondary antibodies (WB 1:10,000; 111-005-144) were from Jackson lab.

### Cell lines, plasmid, and transfection

HeLa cells (ATCC RR-B51S) were routinely grown at 37 ℃ in Dulbecco’s modified essential media (DMEM, Sigma-Aldrich, D6429), supplemented with 10% FBS (Sigma, F7524), 100 U/ml Penicillin/Streptomycin, 2 mM L-Glutamine (L-Gln). Triple *(optn*/*ndp52*/*tax1bp1)* and penta (*tax1bp1*/*optn*/*ndp52*/*nbr1*/*p62)* knockout (KO) HeLa cell lines were kindly provided by Prof. Richard Youle (NINDS, NIH, USA). The individual knockouts of *taxbp1*, *sqstm1/p62* and *sec22b* in HeLa cell lines were generated by CRISPR-Cas9 approach. We used the following targeting sequence to design guide RNAs:

SEC22B: 5’ – CACCGGAGTGCAAATCTTCTAGGT – 3’.

TAX1BP1: 5’ – CACCGTAGTGGTGACCACAAAAGC – 3’,

SQSTM1/p62: 5’ – CACCGCCGAATCTACATTAAAGAGAA – 3’ and

The guide RNA oligonucleotides were annealed and ligated to BpiI digested pSp-Cas9-BB-2A-Puro (PX459) V 2.0 vector (a gift from Feng Zhang; Addgene Plasmid #62988). Cells were transfected with respective plasmids using Lipofectamine-3000 (Thermo Fischer, L3000001) according to the manufacturer’s instructions followed by selection with Puromycin (2-3 μg/ml). The individual clones obtained after selection were expanded to screen them for the expression by immunoblotting using specific antibodies.

Plasmids encoding MYC-tagged UBB^+1^ was described previously (31). SQSTM1/p62 with C-terminal HA tag (HA-SQSTM1/p62, 452 residues) was obtained from addgene (28027, deposited by Qing Zhong). The coding sequence of the SQSTM1/p62 without PB1 (HA-SQSTM1/p62ΔPB1, 330 residues) domain was subcloned from the full-length HA-tagged SQSTM1/p62 plasmid.

PCR-based site-directed mutagenesis (SDM) was used to generate specific point mutations in the full-length HA-tagged SQSTM1/p62 plasmid. To disrupt the LC3-interacting region (LIR), tryptophan at position 340 and leucine at position 343 were substituted with alanine, generating the HA-SQSTM1/p62 W340A/L343A mutant. The primer pair used for this mutation was:

**Forward**: 5′-GGAGATGATGACGCGACCCATGCGTCTTCAAAAG-3′ and

**Reverse**: 5′-CTTTTGAAGACGCATGGGTCGCGTCATCATCTCC-3′.

Additionally, to disrupt the ubiquitin-binding function of the UBA domain, methionine at position 404 was substituted with valine (M404V) in the same plasmid using the following primer pair:

**Forward**: 5′-CAGATGCTGTCCGTGGGCTTCTCTG-3′ and

**Reverse**: 5′-CAGAGAAGCCCACGGACAGCATCTG-3′.

All site-directed mutagenesis reactions were performed using high-fidelity polymerase and verified by DNA sequencing.

### Protein secretion experiments

To measure UBB⁺¹ secretion, cells (1 × 10⁵) were seeded and cultured to ∼70–80% confluency. Cells were transfected with MYC-UBB⁺¹ along with the indicated control or effector plasmids. After 24 h, the culture medium was replaced with 1.5 ml of fresh complete DMEM. Cells were incubated for an additional 16 h, and the conditioned media were then collected. Media were sequentially centrifuged, first at 1,000 × g for 5 min to remove detached cells, and then at 10,000 × g for 30 min to eliminate debris. Equal volumes of cleared media and corresponding whole-cell lysates (WCL) were resolved by SDS-PAGE and analyzed by immunoblotting. Secretion efficiency was assessed by normalizing the UBB⁺¹ signal in the media to its expression level in the WCL.

### Immunoblotting and immunoprecipitation

For immunoblotting and co-immunoprecipitation, cells harvested 24 h or 48 h post transfection were lysed in NP40 lysis buffer (50 mM Tris-HCl pH 7.6, 150 mM NaCl, 5 mM MgCl2, 1 mM EDTA, 1 mM DTT, 1% NP-40, 5% glycerol) supplemented with protease and deubiquitinase inhibitors (20 mM Iodoacetamide). Protein concentration was measured using Bradford reagent (Thermo Scientific). The insoluble fraction consisting of the remaining pellet was washed once with NP-40 lysis buffer, spun at 17,000 x g for 10 min at 4℃, then lysed in 1 X SDS sample buffer and heated to 95℃ for 10 min. Immunoblotting was performed according to a previously described procedure (31, 64).

For co-immunoprecipitation of overexpressed MYC-UBB^+1^ and HA-p62 variants, Cells were co-transfected with the indicated plasmids for 24 h. Subsequently, they were washed three times with ice-cold PBS (Sigma, D8537) and lysed in 500 μl of ice-cold NP40 lysis buffer. Cell lysates were incubated for 2 h with anti-MYC magnetic beads (Bimake, B26301) and anti-HA magnetic beads (MedChemExpress, HY-K0201). Precipitated complexes were washed with lysis buffer and eluted with MYC and HA peptides (GenScript, RP11731 and RP11735S), respectively at a concentration of 400 μg/ml.

For co-immunoprecipitation of endogenous SQSTM1/p62, the cell lysates were incubated for 2–4 h with specific antibody followed by 1–2 h with protein-A/G plus coupled Sepharose beads (GE Healthcare, 17–0885-01). For control experiments, lysates were also incubated with protein A/G beads alone (without antibody) to assess non-specific binding to the beads. To resolve protein complexes for denaturing immunoprecipitation, cell pellets were lysed in 50 μl of ice-cold NP40 lysis buffer supplemented with 1% SDS. Immunocomplexes were washed with NP-40 lysis buffer and elution was performed by boiling for 5 min in 2x SDS-PAGE loading buffer. Equal amounts of proteins were fractionated in a SurePAGE^TM^, Bis-Tris 4–20% gradient gels (GenScript, M00655). After transferring to nitrocellulose membranes (Millipore), the blots were blocked in Tris-buffered saline containing 5% non-fat milk and 0.1% Tween- 20 and incubated with primary antibodies either for 1 h at room temperature or overnight at 4°C followed by incubation for 1 h with the appropriate horseradish peroxidase-conjugated secondary antibodies. Western blot images were visualized by enhanced chemiluminescence (ECL).

### Immunofluorescence and confocal microscopy

HeLa cells were grown to semi-confluency on glass cover slips and transfected with the indicated plasmids using the Lipofectamine 3000 kit as recommended by the manufacturers. After treatment for the indicated time the cells were fixed in 4% paraformaldehyde (Merck, 100496) and permeabilized with 0.1% Triton X-100 in phosphate buffer saline (PBS) for 5 min RT. Blocking in 5% bovine serum albumin (BSA) (Sigma, A7906) in PBS for 30 min at RT was performed before antibody labeling. The cells were labeled in 5% BSA-PBS using rabbit anti-UBB^+1^, mouse anti-HA, mouse anti-SQSTM1/p62, mouse anti-MYC and rabbit anti-LC3 followed by the appropriate Alexa Fluor 488 or 594 conjugated secondary antibodies. The nuclei were stained with 2.5 μg/ml DAPI (Sigma, D9542) in PBS for 10 min. The coverslips were again washed with PBS thrice and finally mounted with Fluoromount G (Thermo) on glass slides and stored at −20 °C until image acquisition. Images were acquired using Zen blue software on an LSM 880 confocal microscope system (ZEISS) with a Plan Apochromat X63 1.4 numerical aperture objective and oil immersion. The colocalization analysis was performed by using the Profile analysis tool in the ZEN lite (V 3.0) software (Zeiss) by fluorescence intensity line measurement approach.

### Gene silencing by RNA interference

All gene specific DsiRNA oligonucleotides were purchased from Integrated DNA Technologies. DsiRNA was transfected with lipofectamine-3000 transfection reagent (Thermo Fischer, L3000001) at a final concentration of 25 nM. Following are DsiRNA constructs used in this study (**Table 1**),

**Table 1.**
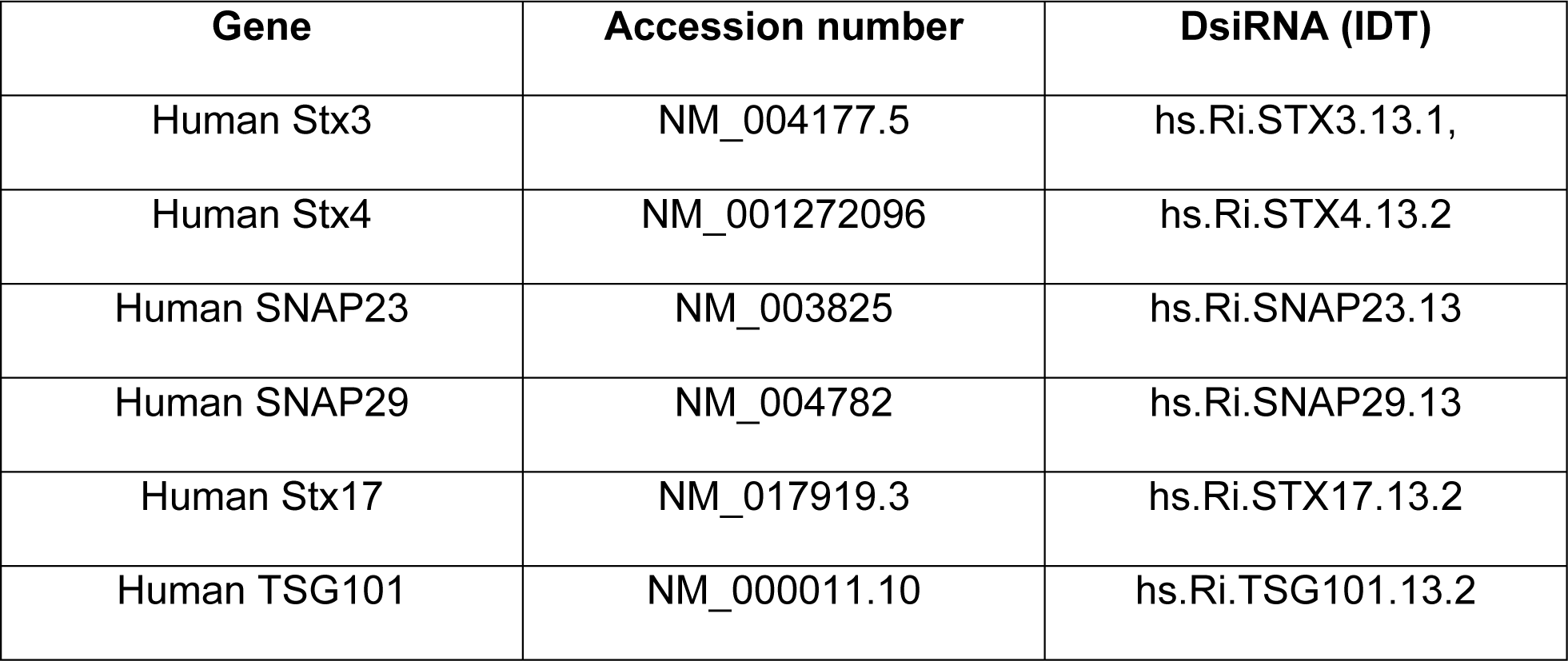
DsiRNA constructs used in this study.

| Gene | Accession number | DsiRNA (IDT) |
| --- | --- | --- |
| Human Stx3 | NM_004177.5 | hs.Ri.STX3.13.1, |
| Human Stx4 | NM_001272096 | hs.Ri.STX4.13.2 |
| Human SNAP23 | NM_003825 | hs.Ri.SNAP23.13 |
| Human SNAP29 | NM_004782 | hs.Ri.SNAP29.13 |
| Human Stx17 | NM_017919.3 | hs.Ri.STX17.13.2 |
| Human TSG101 | NM_000011.10 | hs.Ri.TSG101.13.2 |

| **Gene** | **Accession number** | **DsiRNA (IDT)** |
| --- | --- | --- |
| Human Stx3 | NM_004177.5 | hs.Ri.STX3.13.1, |
| Human Stx4 | NM_001272096 | hs.Ri.STX4.13.2 |
| Human SNAP23 | NM_003825 | hs.Ri.SNAP23.13 |
| Human SNAP29 | NM_004782 | hs.Ri.SNAP29.13 |
| Human Stx17 | NM_017919.3 | hs.Ri.STX17.13.2 |
| Human TSG101 | NM_000011.10 | hs.Ri.TSG101.13.2 |

### Statistical analysis

All statistical analyses were performed using GraphPad Prism (version 5.0). Data are presented as mean ± standard deviation (SD) as indicated in figure legends. For comparisons between two groups, an unpaired two-tailed Student’s t-test was used. For multiple group comparisons, such as the quantification of p62 condensate volume and number per cell, statistical significance was assessed using one-way ANOVA followed by Tukey’s multiple comparisons test. A p-value > 0.05 was considered not significant (ns). Statistical significance is indicated as follows: *p < 0.05, **p < 0.01, and ***p < 0.001.

### AlphaFold-based modeling of UBB⁺¹–p62 UBA interaction

Protein sequences of full length UBB⁺¹ and the UBA domain of human SQSTM1/p62 (residues 387–436) were submitted to the AlphaFold3-based ColabFold pipeline (65) for structural prediction of a protein-protein complex. The top-ranked predicted model (based on the predicted alignment error and pLDDT scores) was selected for visualization. Structural images were rendered using PyMOL (v2.5.4) Color coding was applied as follows: UBB⁺¹ in green, its C-terminal tail in red, the p62 UBA domain in magenta, Met404 in orange, and the hydrophobic patch residues of UBB⁺¹ (Ile44, Val70, and Leu8) in blue. Representative structural snapshots were rendered and included as **Supplementary Fig. S5D.**

## Supporting information

p62 manuscript Supplementary figures S1_S5

## Acknowledgments

We sincerely thank Prof. Richard Youle and his team for kindly providing us with the penta knockout (5KO) and triple knockout (TKO) HeLa cell lines. We thank Dr. Nitsan Dahan and all members of the Microscopy and sorting units of the LS&E institute at the Technion. The authors thank Dr. Inbal Maniv, Noa Reis, and all the Glickman lab members for their intellectual input and critical feedback on the manuscript.

## Abbreviations

AD: Alzheimer disease
CHX: cycloheximide
DsiRNAs: dicer-substrate small interfering RNA
KO: knockout
5 KO: penta knockout
TKO: triple knockout LC3, microtubule associated protein 1 light chain 3
SNARE: SNAP soluble NSF attachment protein receptor
SQSTM1/p62: sequestosome 1
STX: syntaxin
TAX1BP1: Tax1 binding protein 1
WB: Western blot
WT: wild type

## Data Sharing

All relevant data is included with the submission.

## Author contributions

ARW conducted all experiments. MHG obtained funding and supervised. Both authors wrote the manuscript.

