## Supplementary material for "Molecular mechanisms underlying p62-dependent secretion of the Alzheimer-associated ubiquitin variant, UBB^+1^": p62 manuscript Supplementary figures S1_S5

This file contains supplemental figures S1-S5

### Figure S1

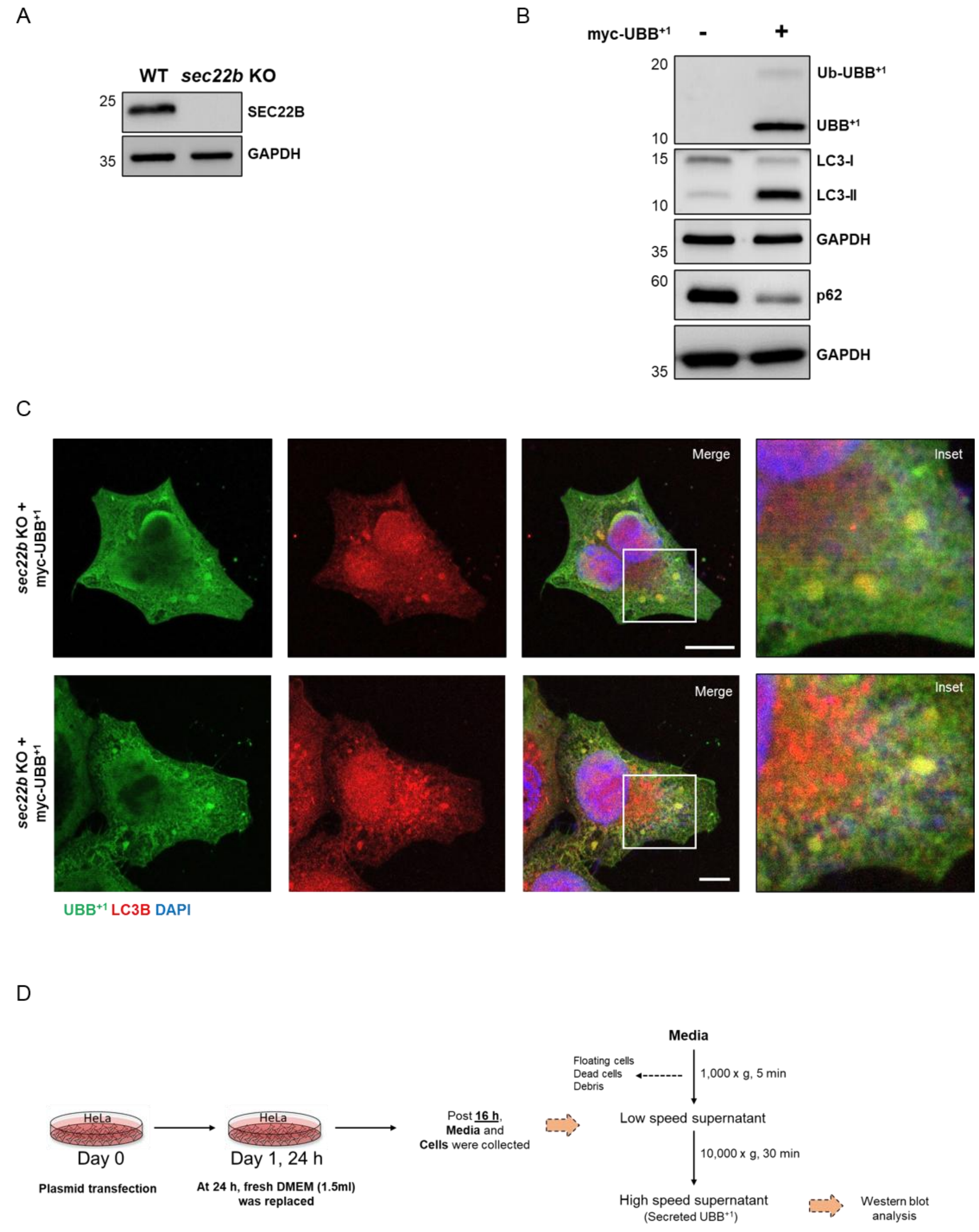
A
WT
sec22b KO
25
SEC22B
35
GAPDH
B
myc-UBB+1
-
+
20
Ub-UBB+1
UBB+1
10
15
LC3-I
LC3-II
10
GAPDH
35
60
p62
GAPDH
35
C
sec22b KO + myc-UBB+1
Merge
Inset
sec22b KO + myc-UBB+1
Merge
Inset
UBB+1 LC3B DAPI
D
Media
Floating cells
Dead cells
Debris
1,000 x g, 5 min
HeLa
Day 0
Plasmid transfection
HeLa
Day 1, 24 h
At 24 h, fresh DMEM (1.5ml) was replaced
Post 16 h, Media and Cells were collected
Low speed supernatant
10,000 x g, 30 min
High speed supernatant
(Secreted UBB+1)
Western blot analysis

### Figure S1

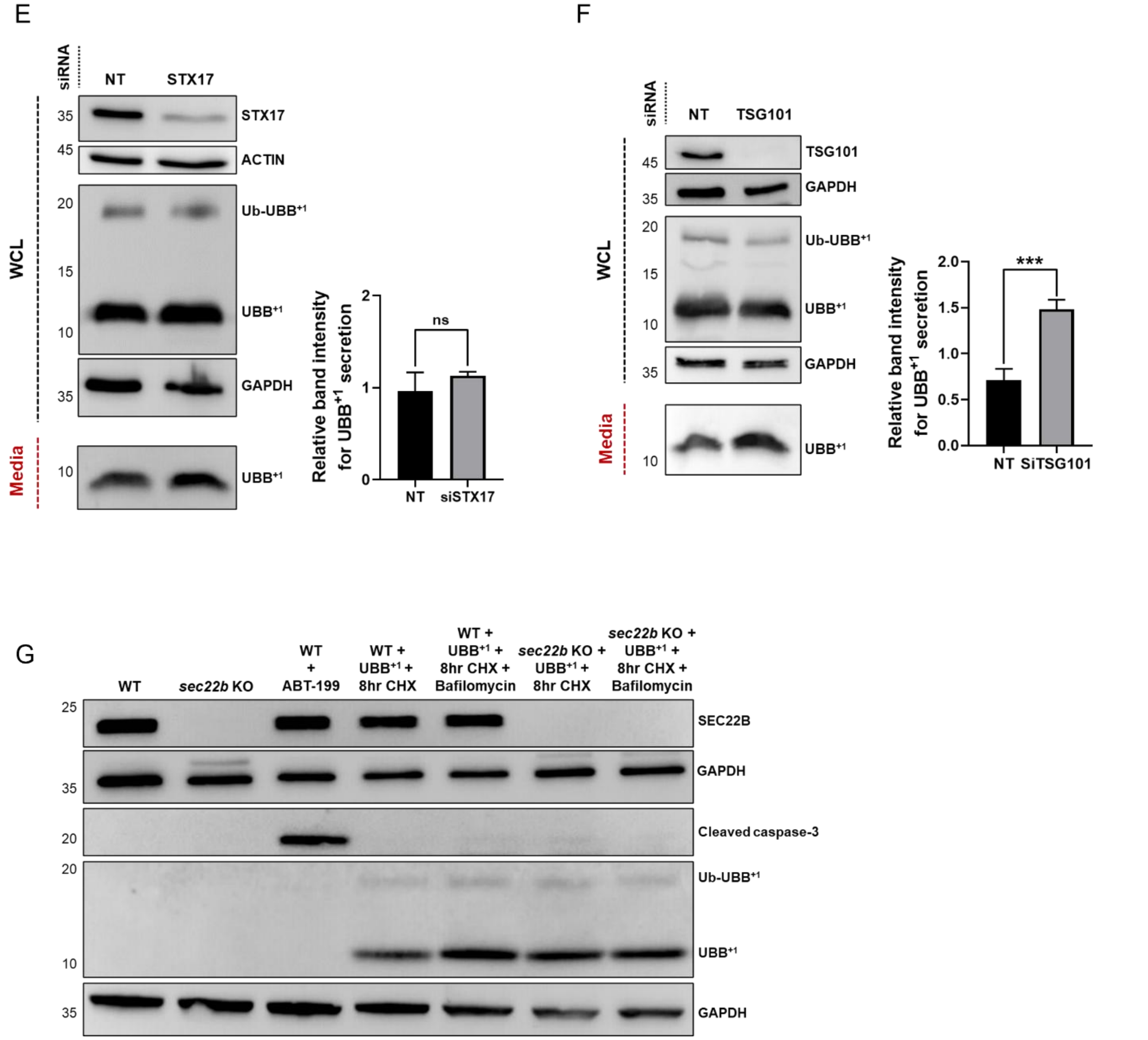
E
siRNA
NT
STX17
WCL
Media
35
45
20
15
10
35
10
STX17
ACTIN
Ub-UBB+1
UBB+1
GAPDH
UBB+1
Relative band intensity for UBB+1 secretion
ns
2
1
0
NT
siSTX17
F
siRNA
NT
TSG101
WCL
Media
45
35
20
15
10
35
10
TSG101
GAPDH
Ub-UBB+1
UBB+1
GAPDH
UBB+1
Relative band intensity for UBB+1 secretion
***
2.0
1.5
1.0
0.5
0.0
NT
SiTSG101
G
WT
sec22b KO
WT + ABT-199
WT + UBB+1 + 8hr CHX
WT + UBB+1 + 8hr CHX + Bafilomycin
sec22b KO + UBB+1 + 8hr CHX
sec22b KO + UBB+1 + 8hr CHX + Bafilomycin
25
35
20
20
10
35
SEC22B
GAPDH
Cleaved caspase-3
Ub-UBB+1
UBB+1
GAPDH

**Figure S1.**

**(A)** Validation of SEC22B knockout in HeLa cells. Immunoblot analysis was performed using anti-SEC22B antibody. GAPDH served as the loading control.

**(B)** Immunoblot analysis of whole-cell lysates from HeLa cells transfected with or without MYC-$UBB^{+1}$. Cells were harvested after 24 h and probed with antibodies against LC3B, $UBB^{+1}$, p62/SQSTM1, and GAPDH. Expression of $UBB^{+1}$ led to an increase in LC3B-II levels and a decrease in endogenous p62 levels, indicative of autophagy induction, consistent with literature reports on autophagosome formation and cargo engagement.

**(C)** Immunofluorescence staining of SEC22B KO HeLa cells expressing MYC-$UBB^{+1}$. Cells were fixed and stained with specific antibodies against $UBB^{+1}$ (green) and LC3B (red), and nuclei were counterstained with DAPI (blue). Merged images and zoomed insets clearly demonstrate colocalization of $UBB^{+1}$ with LC3-positive autophagosomes. Scale bar: 2 μm.

**(D)** Schematic diagram outlining the experimental workflow for the $UBB^{+1}$ secretion assay via differential centrifugation. At 24 h post-transfection, conditioned media were replaced and collected after the indicated duration. Centrifugation speeds and times are noted; supernatants were retained while pellets containing cells, dead cells, or debris were discarded at each step.

**(E–F)** HeLa cells were transfected with DsiRNAs targeting STX17 or TSG101, or a non-targeting (NT) control. Forty-eight hours post-transfection, cells were incubated with fresh DMEM for 16 h. Conditioned media and whole-cell lysates (WCL) were subjected to SDS-PAGE followed by immunoblotting for $UBB^{+1}$. GAPDH and ACTIN were used as loading controls for WCL. Quantification of $UBB^{+1}$ secretion was performed across three biological replicates. Data are presented as mean ± SD. Statistical significance was assessed by one-way ANOVA followed by Tukey's multiple comparisons test; *, **, *** indicate $p < 0.05$, 0.01, and 0.001, respectively; ns = non-significant.

**(G)** Assessment of apoptosis in WT and SEC22B KO HeLa cells following cycloheximide (CHX) and Bafilomycin A1 treatment. Cells were treated with 100 µg/mL CHX alone or in combination with 100 nM Bafilomycin A1 for 0 or 8 h. Whole-cell lysates were immunoblotted for cleaved caspase-3. WT cells treated with the BCL-2 inhibitor ABT-199 (10 µM, 8 h) were used as a positive control for apoptosis induction. No significant cleavage of caspase-3 was observed under CHX or CHX+Bafilomycin conditions, confirming absence of apoptosis.

### Figure S2

A

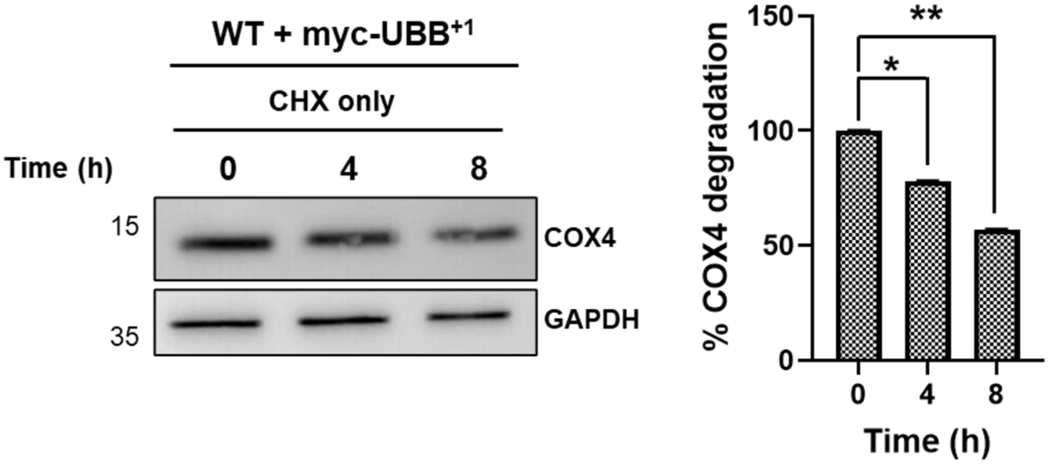
WT + myc-UBB+1
CHX only
Time (h)
0
4
8
15
COX4
35
GAPDH
150
100
50
0
% COX4 degradation
**
*
0
4
8
Time (h)

B

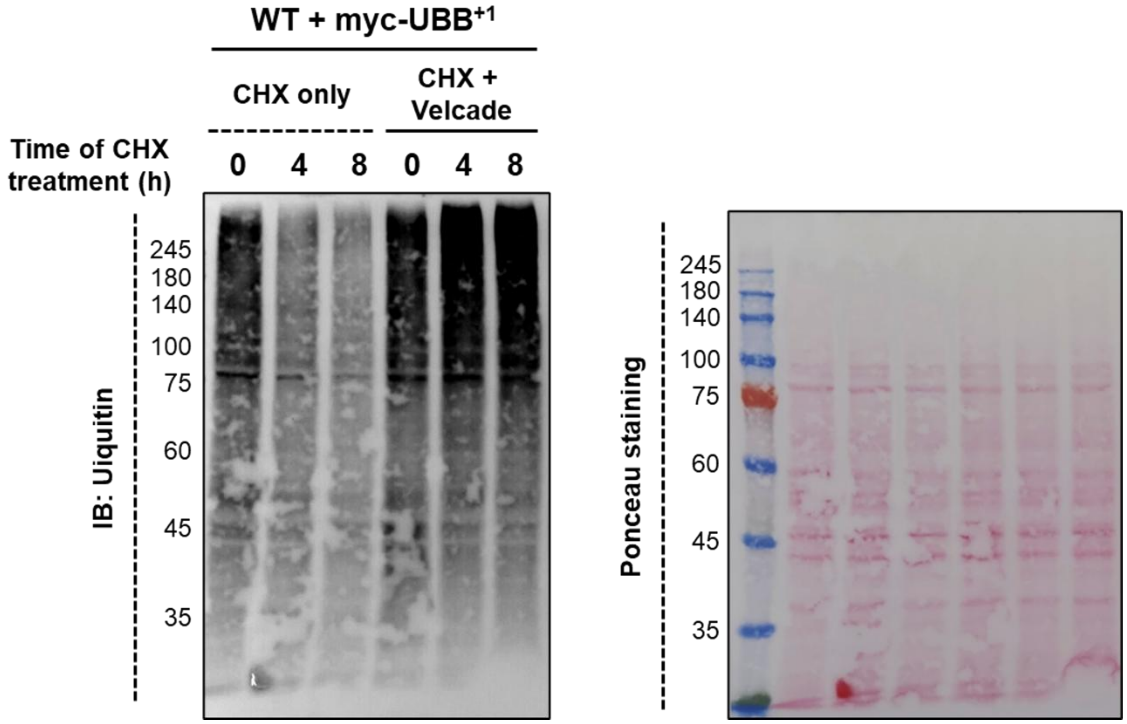
WT + myc-UBB+1
CHX only
CHX + Velcade
Time of CHX treatment (h)
0 4 8 0 4 8
245
180
140
100
75
60
45
35
IB: Uiquitin
Ponceau staining

**Figure S2.**

**(A)** Wild-type (WT) HeLa cells expressing MYC-$UBB^{+1}$ were treated with 100 µg/mL cycloheximide (CHX) for the indicated time points. Whole-cell lysates (WCL) were harvested and subjected to immunoblotting using an anti-COX4 antibody. GAPDH served as a loading control. Representative immunoblots are shown alongside densitometric quantification of COX4 normalized to GAPDH levels. Data represent mean ± SD from three independent experiments.

**(B)** WT HeLa cells expressing MYC-$UBB^{+1}$ were treated with CHX (100 µg/mL) in the presence or absence of the proteasome inhibitor Velcade (100 nM) for the indicated durations. WCL were collected and immunoblotted with an anti-ubiquitin antibody. Ponceau S staining of the membrane is shown as a loading control. Representative blots and quantification from three independent experiments are provided.

For all panels, data are presented as mean ± SD. Statistical significance was determined using a paired two-tailed t-test; *, **, *** indicate $p < 0.05$, 0.01, and 0.001, respectively.

### Figure S3

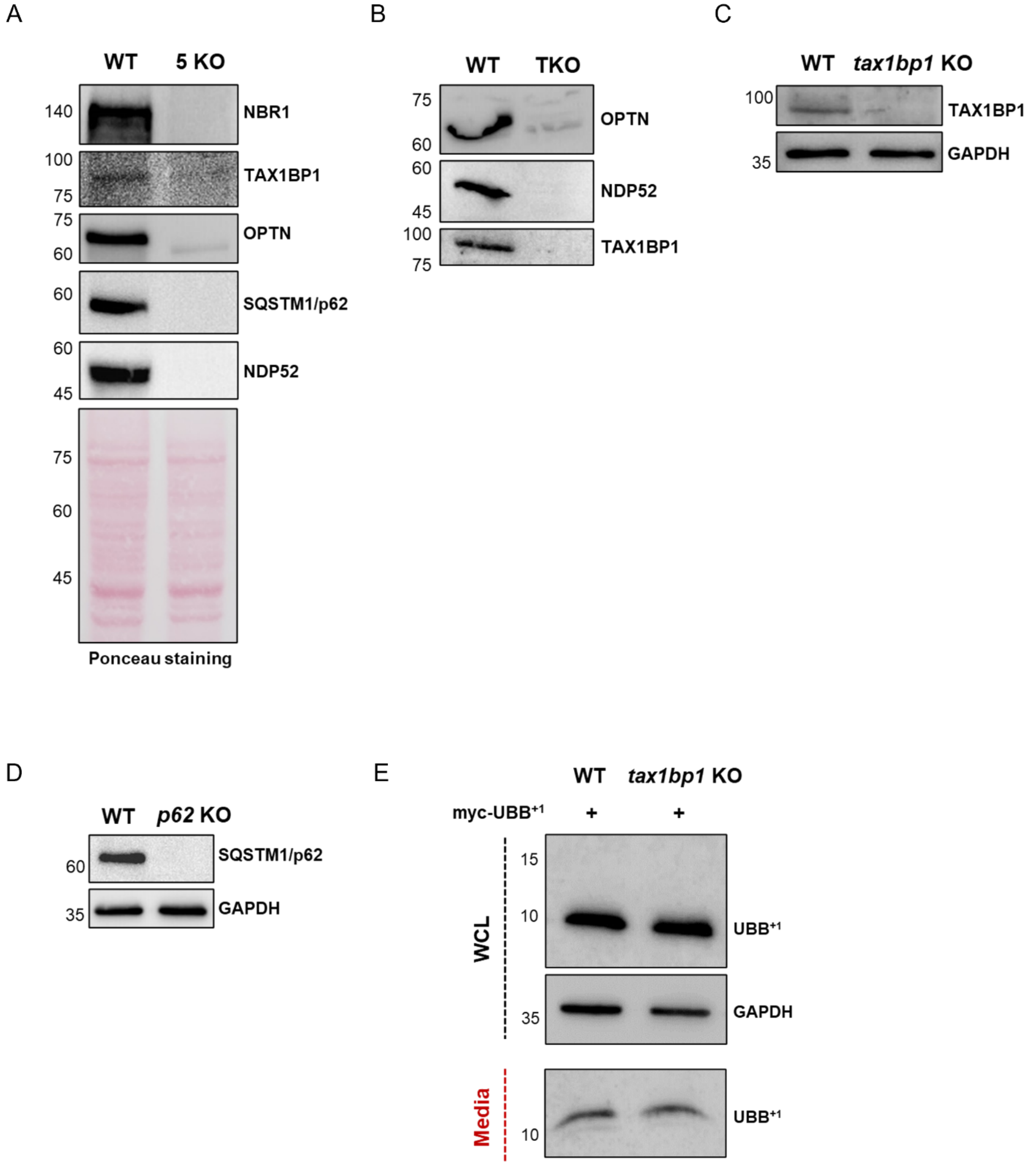
A
WT
5 KO
140
NBR1
100
75
TAX1BP1
75
60
OPTN
60
SQSTM1/p62
60
45
NDP52
75
60
45
Ponceau staining
B
WT
TKO
75
60
OPTN
60
45
NDP52
100
75
TAX1BP1
C
WT
tax1bp1 KO
100
TAX1BP1
35
GAPDH
D
WT
p62 KO
60
SQSTM1/p62
35
GAPDH
E
WT
tax1bp1 KO
myc-UBB+1
+
+
WCL
15
10
UBB+1
35
GAPDH
Media
10
UBB+1

### Figure S3

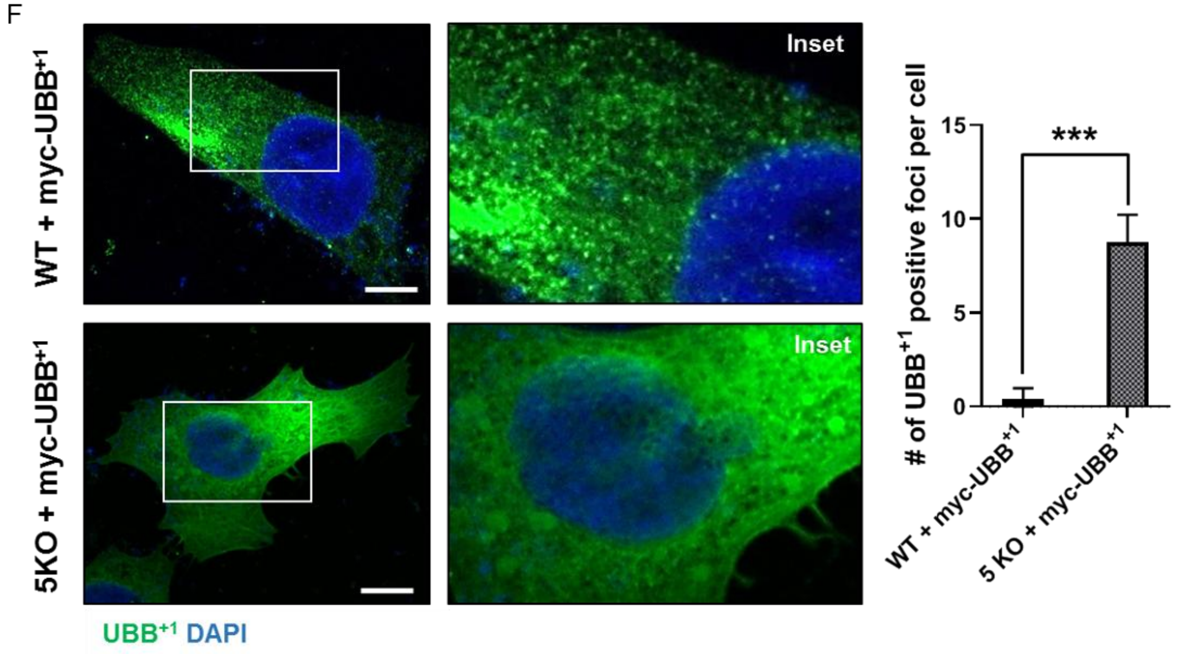
F
WT + myc-UBB+1
5KO + myc-UBB+1
Inset
Inset
UBB+1 DAPI
### of UBB+1 positive foci per cell
15
10
5
0
***
WT + myc-UBB+1
5 KO + myc-UBB+1

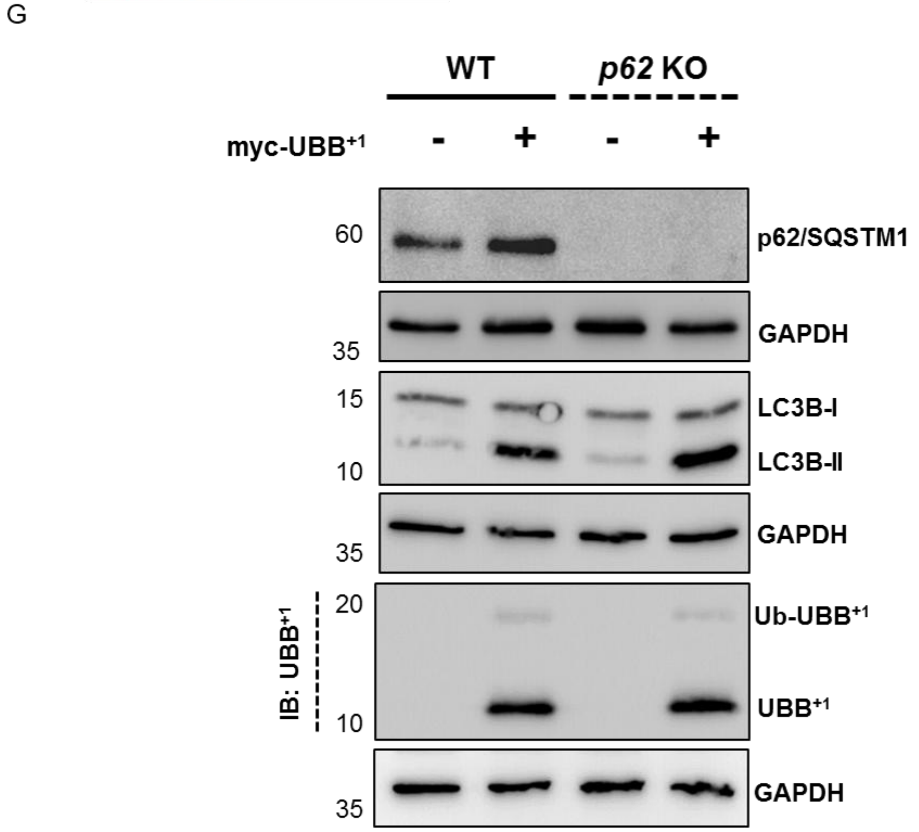
G
WT
p62 KO
myc-UBB+1
-
+
-
+
60
p62/SQSTM1
GAPDH
35
15
LC3B-I
LC3B-II
10
GAPDH
35
IB: UBB+1
20
Ub-UBB+1
UBB+1
10
GAPDH
35

**Figure S3.**

**(A)** Protein levels of SQSTM1/p62, NBR1, OPTN, NDP52, and TAX1BP1 were assessed in WT and 5KO (*tax1bp1*/*optn*/*ndp52*/*nbr1*/*p62)* HeLa cells by immunoblotting. Ponceau staining was used as the loading control.

**(B)** Protein levels of OPTN, NDP52, and TAX1BP1 were determined in TKO (*tax1bp1*/*optn*/*ndp52*/*nbr1*/*p62)* HeLa cells by immunoblotting with specific antibodies.

**(C)** Immunoblotting was performed to analyze the protein levels of TAX1BP1 in *tax1bp1* KO HeLa cells. GAPDH was used as the loading control.

**(D)** p62 expression in *p62* KO HeLa cells was analyzed by immunoblotting. GAPDH was used as the loading control.

**(E)** Conditioned media and WCL from WT and *tax1bp1* KO HeLa cells transfected with MYC-UBB$^{+1}$ were analyzed by immunoblotting after 24 h of transfection and 16 h of incubation in fresh DMEM. GAPDH was used as the loading control for WCL.

**(F)** Immunofluorescence staining of MYC–UBB$^{+1}$ in WT and penta KO (TAX1BP1/OPTN/NDP52/NBR1/p62) HeLa cells. Cells were fixed and stained with anti–UBB$^{+1}$ antibody (green) and counterstained with DAPI (blue) to visualize nuclei. Representative images are shown. In *penta* KO cells, UBB$^{+1}$ displayed a diffuse cytoplasmic distribution with multiple puncta, similar to the pattern observed in *p62* KO cells. This suggests that SQSTM1/p62 loss alone is sufficient to disrupt the subcellular organization of UBB$^{+1}$ aggregates. Scale bar: 2 μm. Statistical significance was determined by a paired t-test. Data represent the mean ± SD of the percentage of cells containing $UBB^{+1}$ foci, assessed in approximately 100 cells. Error bars indicate, means ± SD. For all statistical tests, *, **, ***, $p < 0.05$, 0.01, and 0.001, respectively.

**(G)** Wild-type (WT) and *p62* knockout (KO) HeLa cells were transiently transfected with a MYC-$UBB^{+1}$ expressing plasmid or treated with transfection reagent alone (mock control). After 24 h, whole-cell lysates (WCL) were collected, resolved by 15% SDS-PAGE, and immunoblotted with specific antibodies. LC3-II was detected to assess autophagy induction. GAPDH was used as a loading control.

### Figure S4

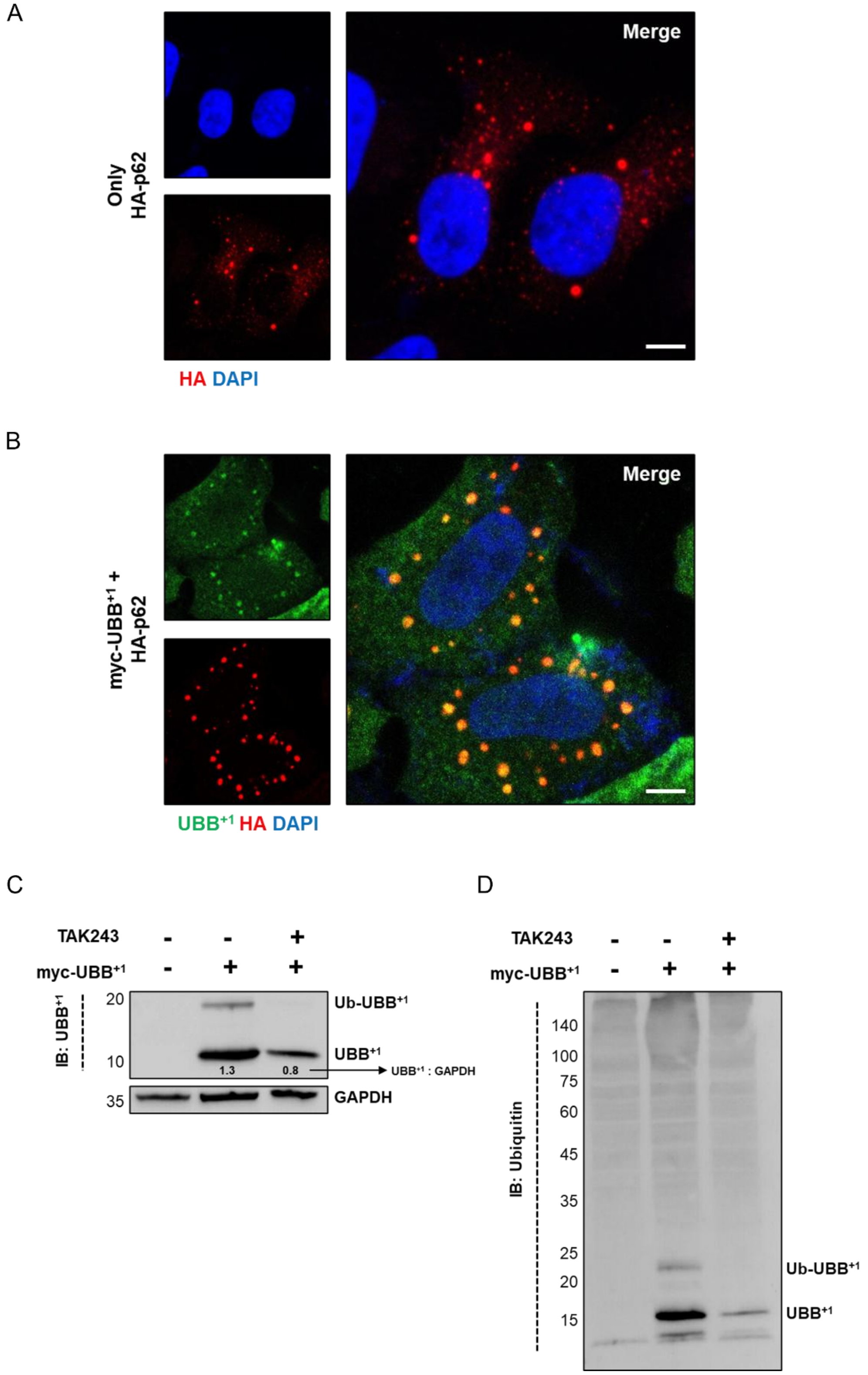
A
Merge
Only
HA-p62
HA DAPI
B
Merge
myc-UBB+1 +
HA-p62
UBB+1 HA DAPI
C
TAK243
myc-UBB+1
IB: UBB+1
20
10
35
Ub-UBB+1
UBB+1
1.3
0.8
UBB+1 : GAPDH
GAPDH
D
TAK243
myc-UBB+1
IB: Ubiquitin
140
100
75
60
45
35
25
20
15
Ub-UBB+1
UBB+1

**Figure S4.**

(**A**) HeLa cells expressing HA-p62 plasmid were fixed and stained for SQSTM1/p62 using HA-specific antibody (red). Scale bar: 2 μm.

(**B**) Immunofluorescence staining of overexpressed HA-p62 in HeLa cells co-expressing MYC-UBB$^{+1}$. Cells grown in complete media were fixed, permeabilized, and stained with monoclonal antibodies against UBB$^{+1}$ (green) and HA (red). Merged images of green and red channels are shown, with nuclei counterstained using DAPI (blue). Scale bar: 2 μm.

**(C and D)** Immunoblot analysis of WCL from HeLa cells expressing MYC-UBB$^{+1}$, treated or untreated for 3 h with 1 μM TAK243 (an inhibitor of ubiquitin-conjugating enzyme E1). UBB$^{+1}$ and its ubiquitinated form (Ub-UBB$^{+1}$) were detected using monoclonal anti-UBB$^{+1}$ antibody. GAPDH served as a loading control. Quantification of UBB$^{+1}$ band intensity relative to GAPDH is indicated. In parallel, polyubiquitinated proteins were analyzed with an anti-ubiquitin antibody.

### Figure S5

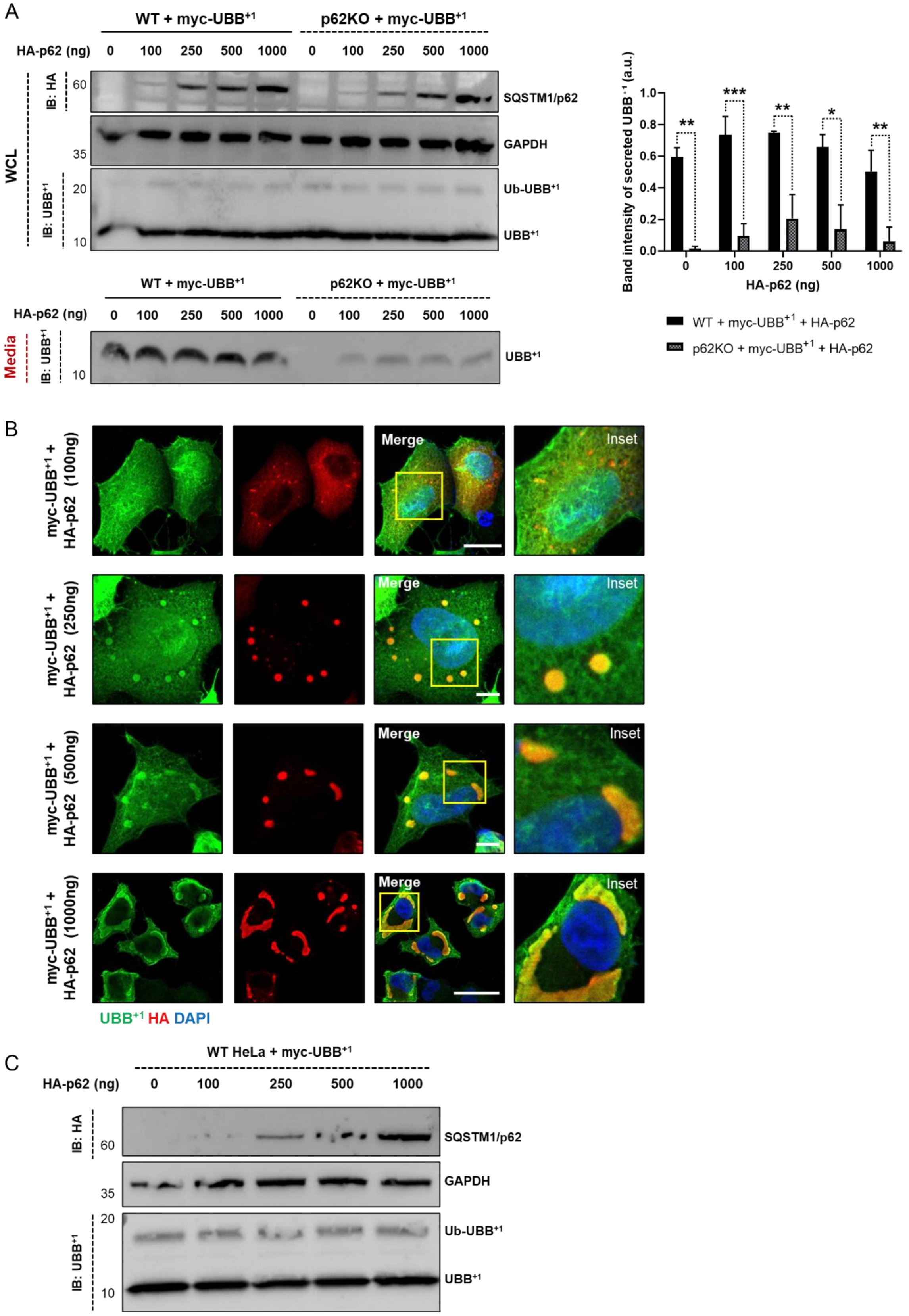
A
WT + myc-UBB+1
p62KO + myc-UBB+1
HA-p62 (ng) 0 100 250 500 1000 0 100 250 500 1000
WCL
IB: HA
60
SQSTM1/p62
GAPDH
35
IB: UBB+1
20
Ub-UBB+1
10
UBB+1
Band intensity of secreted UBB+1 (a.u.)
1.0
0.8
0.6
0.4
0.2
0.0
**
***
**
*
**
HA-p62 (ng)
WT + myc-UBB+1 + HA-p62
p62KO + myc-UBB+1 + HA-p62
Media
B
myc-UBB+1 + HA-p62 (100ng)
myc-UBB+1 + HA-p62 (250ng)
myc-UBB+1 + HA-p62 (500ng)
myc-UBB+1 + HA-p62 (1000ng)
Merge
Inset
UBB+1 HA DAPI
C
WT HeLa + myc-UBB+1
HA-p62 (ng) 0 100 250 500 1000
IB: HA
60
SQSTM1/p62
GAPDH
35
IB: UBB+1
20
Ub-UBB+1
10
UBB+1

### Figure S5

D

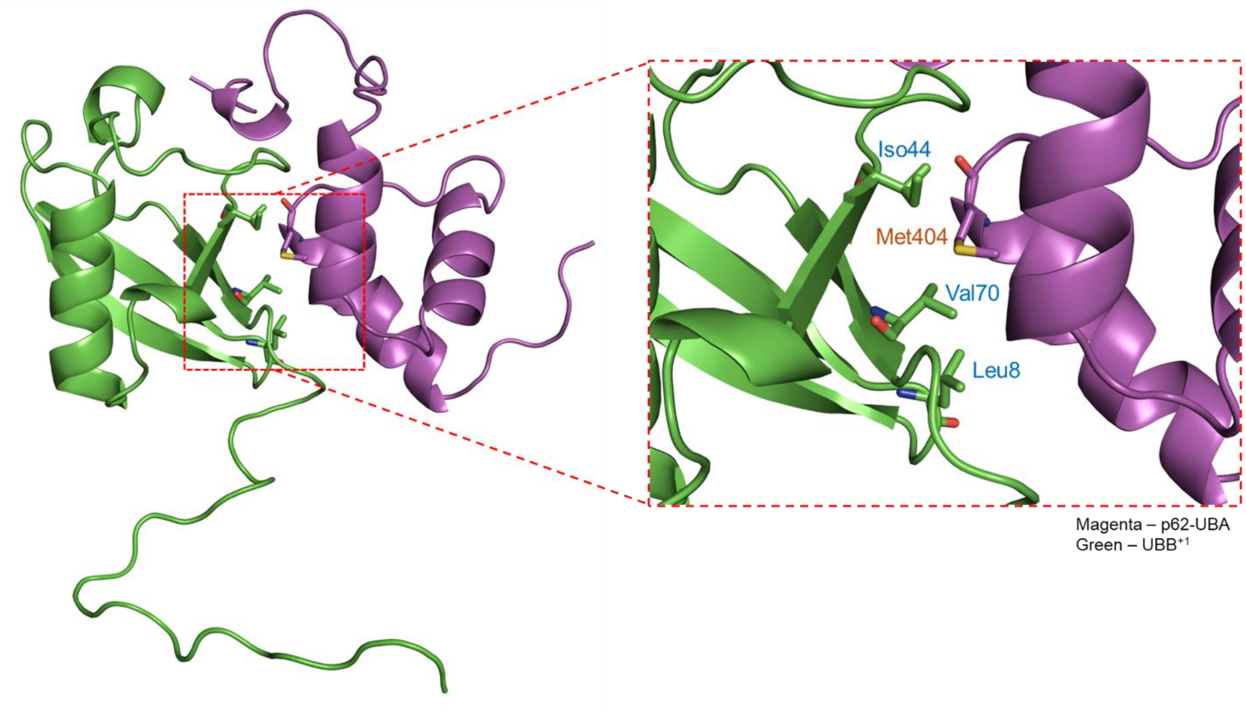

#### Figure S5.

**(A)** WT and p62 knockout (KO) HeLa cells were co-transfected with MYC–$UBB^{+1}$ and increasing concentrations of HA–p62 FL (0 ng, 100 ng, 250 ng, 500 ng, and 1 µg). After 24 h, cells were incubated in fresh DMEM for an additional 16 h, and conditioned media were collected. $UBB^{+1}$ secretion was assessed by immunoblotting. In WT cells, increasing HA–p62 levels resulted in consistent secretion of $UBB^{+1}$. In p62 KO cells, secretion was restored at intermediate plasmid concentrations (250–500 ng), while higher expression (1 µg) led to reduced secretion, possibly due to excessive condensate formation, consistent with observations in panel (B). Equal volumes of media were collected from comparable cell numbers across all conditions.

**(B)** WT HeLa cells were co-transfected with MYC–$UBB^{+1}$ and increasing amounts of HA–p62 FL (0 ng, 100 ng, 250 ng, 500 ng, and 1 µg). After 24 h, cells were fixed and stained with antibodies against HA (red) and $UBB^{+1}$ (green). Nuclei were counterstained with DAPI (blue). Representative immunofluorescence images demonstrate progressive p62 body formation with increasing plasmid concentrations.

**(C)** Corresponding western blot analysis of whole-cell lysates (WCL) from WT HeLa cells transfected as in (A), immunoblotted with antibodies against HA and $UBB^{+1}$ to confirm protein expression. GAPDH was used as a loading control.

**(D)** Representative structural models were visualized using PyMOL based on AlphaFold3 Multimer predictions. $UBB^{+1}$ is shown in green. The SQSTM1/p62 UBA domain is displayed in magenta. A zoomed-in view illustrates the close spatial proximity between

Met404 (colored orange) and the hydrophobic patch residues of $UBB^{+1}$, Ile44, Val70, and Leu8, colored in blue. These residues form a hydrophobic interface predicted to mediate binding between $UBB^{+1}$ and the UBA domain of p62. The model supports experimental data showing loss of interaction upon mutation of Met404 to valine (M404V), which increases the inter-residue distance and disrupts the interaction.
